# Microbial community characteristics and stable relationships centered on anammox bacteria revealed by global-scale analysis

**DOI:** 10.1101/2025.03.21.644508

**Authors:** Yi-Cheng Wang, Hui-Min Fu, Hong-Yuan Liu, Han Wu, Zi-Han Zeng, Jian-Hong He, Meng-Jiao Gao, Peng Yan, Liang Zhang, You-Peng Chen

## Abstract

Anaerobic ammonium-oxidizing bacteria (AnAOB) play important roles in both artificial wastewater treatment systems and natural ecosystems. To date, AnAOB pure cultures are not available and they tend to coexist with various microbial species. However, anammox community characteristics including the relationships between AnAOB and their companion bacteria at the global perspective and their impacts on anammox metabolism remain unclear. Here, we systematically analyzed the characteristics of anammox communities and the stable relationships concerning AnAOB using a global dataset containing 619 anammox-related amplicons. Different anammox systems showed significant differences in alpha and beta diversity, but shared some core taxa of interest. A total of 89 and 55 core genera and species were identified respectively across anammox communities worldwide, which formed the backbone of artificial anammox systems. Through the analysis of co-abundance networks derived from four distinct artificial anammox systems—biofilm, granular sludge, flocculent sludge, and planktonic cells—we identified 208 stable and 92 limited stable relationships associated with AnAOB. Functional analysis suggested that stable positively-correlated companion bacteria may provide essential cofactors (e.g., molybdenum cofactor, tetrahydrofolate, and coenzyme A) to AnAOB. The companion bacteria which showed limited stable positive correlations with AnAOB in the anammox attachment-growth systems, may mutualize with AnAOB via type pili. This study deepens the understanding of anammox communities, anammox core microbiome, and AnAOB symbiotic relationships. These (limited) stable companion bacteria and corresponding cofactors can potentially guide the development and application of bioaugmentation methods, synthetic anammox communities, and deterioration biomarkers for engineered anammox systems.

## Introduction

Anaerobic ammonium oxidation (anammox) is recognized as a representative of energy-efficient wastewater nitrogen removal processes and an important part of the global nitrogen cycle^1,2^. Anammox is mediated by a deeply branching group within phylum Planctomycetota known as anaerobic ammonium-oxidizing bacteria (AnAOB), which can oxidize ammonium using nitrite as an electron acceptor under anoxic conditions^3^. It is well known that AnAOB has metabolic diversity and habitat diversity^4,5^. AnAOB inhabit not only biological wastewater treatment systems, but also a wide range of natural anoxic environments such as oceans, freshwaters, and sediments^6,7^. Despite extensive research efforts, pure cultures of AnAOB remain elusive due to their exceptionally slow growth rates and dependence on diverse species in complex microbial communities^8,9^. Genome-based analyses have further revealed that AnAOB possess incomplete biosynthetic pathways for essential growth factors, strongly indicating their obligate symbiotic relationships with other microorganisms^10,11^. This feature poses a challenge in studying their functions and metabolisms. Moreover, the presence of diverse symbiotic and antagonistic relationships within anammox communities and the sensitivity of AnAOB to environmental perturbations are challenges for the wide engineering application of anammox processes^12–14^. Revealing the general composition, structure, and symbiotic relationships of anammox communities will help to elucidate AnAOB growth and metabolism. This fundamental understanding will enable precise enhancement of AnAOB activity and growth rates in engineered anammox systems, as well as the guide for isolation of pure AnAOB cultures and synthesizing anammox communities with robustness and efficiency.

Previous studies suggest that anammox communities may share a number of core species within Planctomycetota, Proteobacteria, Chloroflexota, Bacteroidota, and Acidobacteriota^14,15^. However, most anammox community studies have relied on limited samples or specific conditions, including various environmental stresses and reactor types^14,16,17^. The current understanding of core species and their symbiotic relationships with AnAOB is largely based on fragmented and system-specific studies, limiting the generalizability of these findings. A systematic and global characterization of anammox communities, particularly the identification of core microbial taxa and their stable interactions with AnAOB, represents a critical knowledge gap in anammox microbial ecology^18^. Recent efforts to catalog anammox microbiomes using 236 metagenomic samples have advanced our knowledge^8^, yet technical limitations in metagenomic sequencing and analysis may overlook low-abundance but functionally important species^19,20^. In addition, numerous studies have constructed the co-abundance network of anammox communities in different systems to analyze network complexity and AnAOB symbiotic relationships^8,21,22^. Shared species that maintain stable relationships despite different perturbations are very likely to be core species with important roles in microbial communities^23^. Nevertheless, the stability of AnAOB symbiotic relationships across different anammox systems remains unclear.

To address these gaps, we conducted a large-scale comparative analysis of 619 high-quality microbial amplicon samples from diverse global sources. This study aims to: (1) characterize community diversity and composition across different anammox systems, (2) identify core taxa and community structures across different anammox systems, and (3) elucidate stable relationships and potential interactions centered on AnAOB across different anammox systems. This study provides fundamental insights into anammox microbial ecology and offer the guidance for precise regulation or construction of engineered anammox systems.

## Methods

### Sample and metadata collection

A total of 94 anammox samples were collected in this study, including 73 granular sludges, 19 planktonic cells, and 2 biofilms. These samples were collected in our lab-scale anammox bioreactors and details are given in **Supplementary Table 1**.

To obtain more data on anammox communities, 16S rRNA amplicon data related to anammox in the National Center for Biotechnology Information (NCBI) Sequence Read Archive (SRA) database were collected based on keyword searches in March 2024. The keywords of the data search were “anammox”, “anaerobic ammonium oxidation”, and “anaerobic ammonia oxidation”. Other amplicon data that might be involved in anammox were also collected, such as sludge, wastewater, seawater, freshwater, sediment, paddy soil. In total, approximately 100,302 amplicon sequencing data were collected from the NCBI SRA database. The following criteria were used to screen suitable amplicon sequencing data: (1) sequencing data were derived from DNA samples; (2) universal 16S rRNA primers were used for amplification; (3) the constructed libraries were sequenced on the Illumina platform using the paired-end strategy; (4) the number of sequences was no less than 10,000; (5) the average length of merged sequences ranged from 280 to 480 bp; (6) AnAOB is the abundant species in microbial community with a relative abundance of no less than 0.5%. Criteria 1 to 5 were designed to minimize the bias introduced by sample types, amplification primers, sequencing strategies, and platforms. Criterion 6 referred previous studies^24,25^. Metadata on these 16S rRNA amplicon sequencing data were collected according to the NCBI SRA data description and the corresponding publications. Sequencing data that lacked essential sample descriptive information (e.g., sample morphology) were removed. Finally, 525 publicly available amplicon sequencing data were collected and used for downstream analysis (**Supplementary Table 2)**. Amplicon data generated from 94 anammox samples collected in this study also met the six screening criteria described above. All high-quality anammox amplicon data were combined with a total number of 619, containing six sample types: biofilm, granular sludge, flocculent sludge, planktonic cells, non-saline sediment, and saline sediment.

### DNA extraction and high–throughput sequencing

Total DNA of the collected anammox sludges was extracted using the MagAttract PowerSoil Pro DNA Kit (QIAGEN, Germany). DNA concentration and purity were checked using Quantus Fluorometer (Promega, USA) and NanoDrop 2000 (Thermo Fisher Scientific, USA), respectively. DNA integrity was checked using agarose gel electrophoresis. The universal primers 338F (5’-ACTCCTACGGGAGGCAGCAG-3’) and 806R (5’-GGACTACHVGGGTWTCTAAT-3’)^26^ was used for 16S rRNA amplification of the qualified DNA samples, and then constructed libraries were sequenced on the Illumina MiSeq platform using the paired-end strategy.

### Data pre-processing and general bioinformatic methods

Fastq-dump v3.0.7 was used for converting the NCBI SRA amplicon data into the fastq format files. Subsequently, all the 619 amplicon sequencing data were processed using Trimmomatic v0.33^27^ to remove low-quality reads and then merged by FLASH v1.2.7^28^.

Although the amplicon data were generated using universal 16S rRNA primers, the target regions were not all the same across samples. Therefore, this study applied a workflow that did not rely on the reference database, i.e., the sequences of all the amplicon data were trimmed, and only the shared V4 region of 16S rRNA was retained for downstream analysis^29,30^. This reference database-independent workflow preserved the integrity of data as much as possible. The specific processes were as follows: (1) each sequence and the V4 region primers 515F (5’-GTGYCAGCMGCCGCGGTAA-3’) and 806R (5’-GGACTACNVGGGTWTCTAAT-3’)^31^ were aligned using BLAST (https://blast.ncbi.nlm.nih.gov/Blast.cgi); (2) sequences were trimmed to the V4 region according to the alignment results of each sequence.

To improve the accuracy of species identification, the amplicon sequence variants (ASVs) analysis workflow was performed. VSEARCH v2.22.1^32^ was used to infer ASVs and generate ASV table. The sequences were first filtered to eliminate low quality (--fastq_maxee_rate 0.01) and abnormal length (--fastq_maxlen 285 --fastq_minlen 215) sequences. Sequence length filtering criteria referred to a public amplicon analysis workflow (https://astrobiomike.github.io/amplicon/dada2_workflow_ex). The filtered sequences were dereplicated and denoised to infer the ASVs. After removing the chimera ASVs by both uchime3_denovo method and the method that rely on reference database of SILVA v138.1 SSU, the ASV table was then generated according to the mapping sequences. The command classify-sklearn of QIIME 2 v2024.5^33^ was used for taxonomic classification of ASVs based on the Greengenes2 v2022.10 database^34^. Singleton ASVs and ASVs that were not assigned at the kingdom rank were removed. ASVs assigned to “Chloroplast”, “Mitochondria”, “Eukaryota” and “Archaea” were also removed, and the percentage of every part was given in **Supplementary Table 3**. Although some species have been renamed in recent studies, the Greengenes2 taxonomic names were adopted for clarity and accuracy of discussion. The phylogenetic trees of the given ASVs were constructed using the QIIME 2 FastTree^35^ method. PICRUSt2^36^ was used to predict the Kyoto Encyclopedia of Genes and Genomes (KEGG) function and contribution of bacteria in anammox communities with default parameters (--stratified --per_sequence_contrib).

### Analysis of community diversity and composition

Unless otherwise noted, all statistical analyses of anammox communities were performed in R software v.4.4.1. The mecodev package v0.2.0^37^ was used for rarefaction curve plotting to determine sampling size. Prior to community diversity analysis, the ASV table was randomly rarefied according to the minimum number of sample sequences (n = 11549). The alpha diversity index of samples was calculated using the microeco package v1.9.1^37^. The Coverage, Observed richness, ACE, and Shannon indexes were chosen to assess the alpha diversity of anammox communities and statistical significance was tested using analysis of variance (ANOVA). Rare ASVs occurring in only one sample were eliminated before calculating beta diversity. The beta diversity of samples was then calculated using the vegan package v2.7.0^38^ based on the Clark distance and non-metric multidimensional scaling (NMDS) ordination methods. Permutational MANOVA (ADONIS) was used to test the significance of differences in beta diversity. Relative abundances of species were calculated based on rarefied ASV table. Unique and shared ASVs were analyzed for different types of samples. In this study, ASVs assigned to Candidatus Brocadiales order were considered to represent AnAOB^3^.

### Identification of core and conditionally rare or abundant taxa

Core taxa and conditionally rare or abundant taxa (CRAT) in anammox communities were defined based on their relative abundance and frequency of occurrence as previously described^39^. Bacterial species (with average relative abundance > 0.1%) that occurred more than 80%, 50%, and 20% in all samples were defined as strict core, general core, and loose core, respectively. CRAT were defined as having a relative abundance greater than 1% in at least one sample. Besides, taxa that were not assigned at the given taxonomic rank were combined as “unclassified”. Except for the five taxa mentioned above, the rest of the taxa were combined into “other”. There was no overlap between these six taxa.

### Analysis of (limited) stable relationship

The co-abundance networks of anammox communities were constructed and analyzed as described below. The natural samples were not included in the analysis of (limited) stable relationship because the anammox communities in the natural and artificial systems were very different and shared almost no AnAOB. Additionally, the possible outlier samples outside the 95% confidence interval of the beta diversity analysis were excluded to avoid causing bias. Four types of artificial anammox systems (biofilm, granular sludge, flocculent sludge, and planktonic cells) were used to construct the network independently. ASVs that occurred less than 20% in samples were eliminated^40^. Since the SparCC algorithm is more suitable for the analysis of sparse compositional data than conventional correlation computation methods^41^, the SparCC correlation and significance between ASVs were calculated using Fastspar v1.0.0^42^ based on 50 iterations and 1000 permutations, respectively. Correlations with P < 0.01 were considered significant and retained for downstream analysis^43^.

Identification of AnAOB stable relationships referred to the methods described by a previous study^23^. Considering the types and number of collected samples, two kinds of stable relationships were defined: stable relationship and limited stable relationship. Any two ASVs with no less than 75% concordance of significant correlations across four independent networks were defined as having a stable relationship, but the simultaneous presence of two opposite significant correlations was not included. Limited stable relationship was defined as any two ASVs having a consistent significant correlation in two independent networks of attached growth mode (biofilm and granular sludge) while having a consistent insignificant correlation in two independent networks of suspended growth mode (flocculent sludge and planktonic cells). If one species of the (limited) stable relationship assigned to AnAOB, the other species was considered to be a (limited) stable correlated companion of AnAOB. The (limited) stable correlation coefficient was defined as the average significant correlation coefficients of the different networks. Correlations with coefficient ≥ 0.5 or ≤ −0.5 and P < 0.01 were considered strong correlations^44^. Cytoscape v3.10.3^45^ was used to visualize the co-abundance networks.

## Results and discussion

### The global samples and dataset for anammox communities

A global-scale anammox dataset with 619 samples were analyzed to gain a comprehensive understanding of anammox communities and AnAOB. Each sample of the anammox dataset was categorized and labeled with three levels of type information Each sample of the curated anammox dataset was categorized and labeled with three levels of type information (**Fig. 1A**). Artificial system samples dominated the collected samples. This may be due to the fact that a stringent sample collecting criterion was applied, i.e., AnAOB must to be the abundant species (relative abundance ≥ 0.5%) within the given community. Typically, AnAOB can be highly enriched in artificial wastewater treatment systems^46,47^. Lab-scale samples accounted for 80.3% of the samples from the artificial system, and the anammox biomass exists in artificial systems presented in the form of biofilm, granular sludge, flocculent sludge, or planktonic cells^17^. Biofilms and granular sludges were the main types of anammox samples. The collected samples were from four continents: Asia, Europe, North America, and South America (**Fig. 1B**). The total number of sequences in the anammox amplicon dataset was 41.7 million, with a total base number of 10.5 Gbp and an average of 67,298 sequences per sample (**Supplementary Table 4**). A 16S rRNA amplicon dataset of anammox communities was constructed based on the global samples and the results of this study. This dataset has been deposited in Figshare and is openly accessible, including all bacterial ASV sequences identified, ASV taxonomy table, ASV count table, rarefied ASV count table, etc.

**Figure 1.**
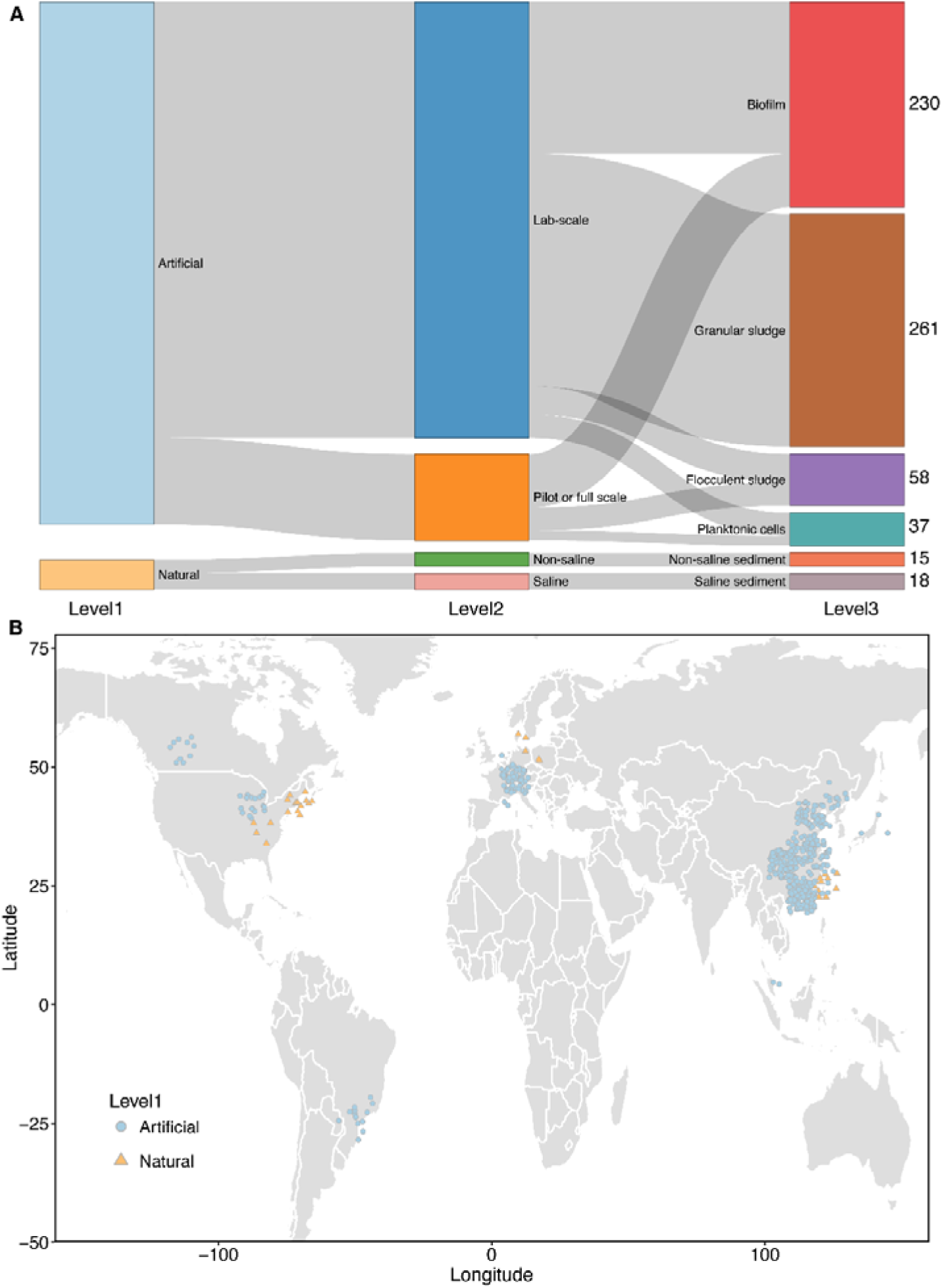
Types and provenance of collected samples. (**A**) The type of samples, including three levels of type information; the number on the right of the figure represents the number of samples in each given level3 type. (**B**) Global distribution of samples. For clarity, the locations of samples were slightly adjusted by the ggplot2 package geom_jitter function. Samples lacking locational information are not shown.

### Alpha and beta diversity of anammox communities

A high-resolution ASV analysis workflow was performed to assess the general diversity characteristics of anammox communities, and results were presented in **Fig. 2**. A total of 30,214 bacterial ASVs were obtained, with an average of 55,642 sequences per sample. The rarefaction curves showed that the Shannon index had leveled off with the number of sampled sequences above 6,000 (**Fig. 2A**). Moreover, the minimum and average Coverage index of 0.93 and 0.98 were calculated after rarefaction, respectively, indicating that the rarefied data could represent the vast majority of bacteria within the anammox communities (**Fig. 2B**).

**Figure 2.**
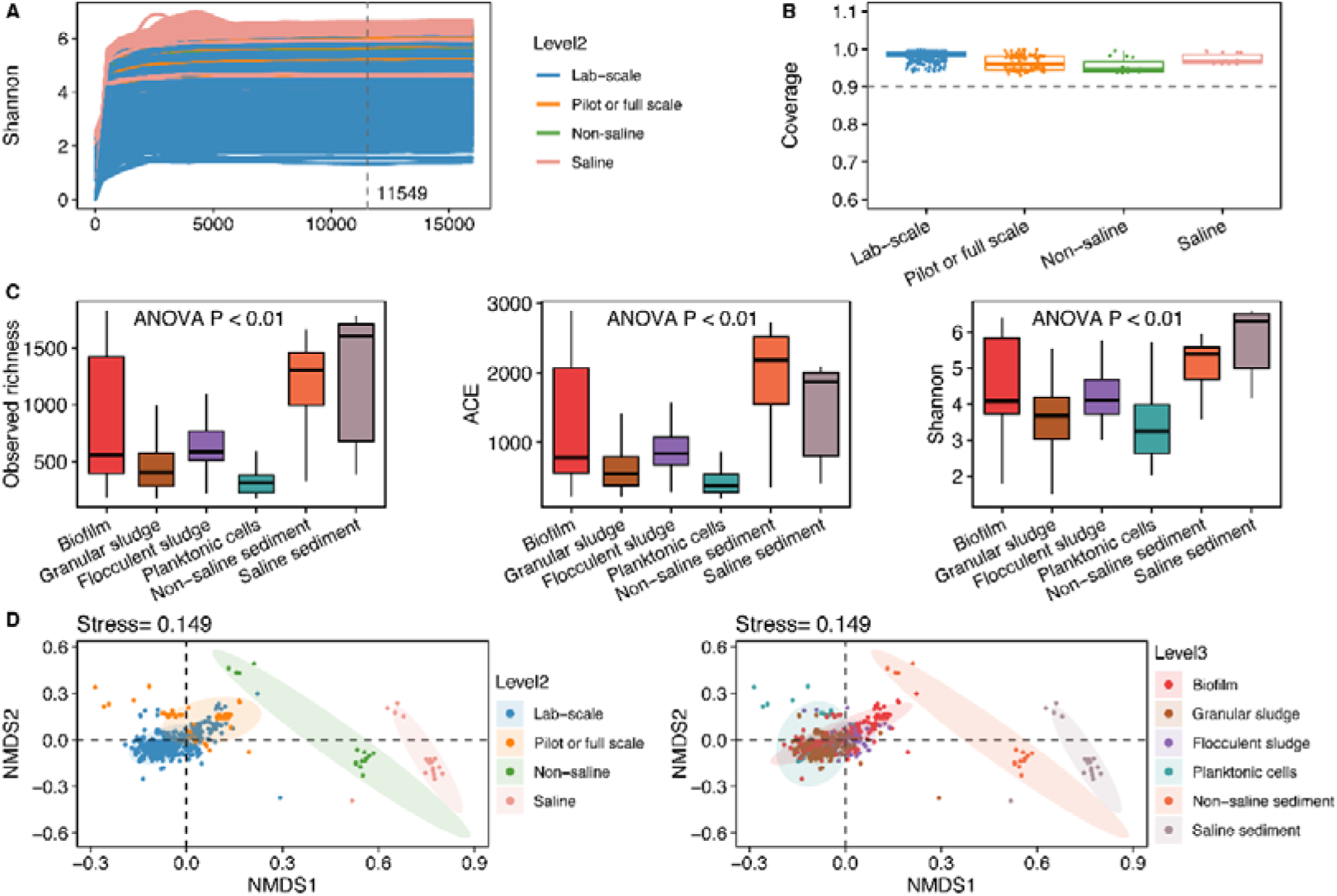
Alpha and beta diversity of global anammox communities. (**A**) Rarefaction curve of level2 samples. (**B**) Coverage index of level2 samples. (**C**) Comparison of three alpha diversity indexes for level3 samples. The difference of alpha index averages between the given types was significant by ANOVA test with P < 0.01. (**D**) Beta diversity of level2 and level3 samples. The difference of beta diversity between the given types was significant by ADONIS test with P < 0.01.

At level3, the alpha diversity of the six types of samples was significantly (P < 0.01) different (**Fig. 2C**). Observed richness, ACE, and Shannon index were all significantly (P < 0.01) higher in the natural samples than in the artificial system samples. This may be attributed to the fact that most of the artificial systems had intentionally adopted AnAOB enrichment strategies, resulting in species diversity loss. Of the four artificial system samples, biofilm had the highest alpha diversity, followed by flocculent sludge and the lowest by planktonic cells. The result suggests that planktonic cell culturing method could be advantageous in obtaining highly enriched AnAOB, which strongly supports the previous findings from a global view^6,47^.

Beta diversity analyses found that the anammox communities of the natural and artificial systems were very distant (**Fig. 2D**), while the two type systems shared only 1,888 (less than 22.9% of the total number of species) species (**Fig. S1A**), suggesting large differences. Despite the proximity of the samples from the artificial system to each other, ADONIS test showed that the community structure was significantly different (P < 0.01) at both level2 and level3, implying that anammox communities assembled in different habitats have different degrees of variation. Only 82 and 43 species were shared by all the different types of systems at the level 2 and level 3, respectively (**Fig. S1B** and **Fig. S1C**). The number of species contained in the four artificial systems varied considerably, with the biofilm system having the highest number of species (n = 16,957) and number of unique species (n = 5,395). This is consistent with the result that biofilms have the highest alpha diversity among artificial systems. But these four artificial systems shared 1,743 species (**Fig. S1D**), and the number of shared species far exceeded the number of species unique to the flocculent sludge and planktonic cell systems. This result implies that there may be some core species in the anammox communities of these artificial systems, and such species are able to adapt to different culture conditions.

### Taxonomy and composition of anammox communities

To explore the general composition and structure of anammox communities, taxonomy classification and relative abundance calculations were performed for all obtained ASVs. The result showed that the percentage of ASV classification at the phylum and genus rank was respectively 96.5% and 62.7% (**Fig. S2A**), with a total of 91 phyla (**Fig. S2B** and **Supplementary Table 5**) and 2699 genera identified. The number of members within Proteobacteria, Chloroflexota, Bacteroidota, Planctomycetota, Firmicutes_A, Actinobacteriota, Myxococcota_A_473307, and Acidobacteriota occupied the top 8, accounting for 71.6% of the total number of ASVs (**Fig. 3**). ASVs assigned to Proteobacteria had a higher overall classification percentage at lower taxonomic ranks than those assigned to other phyla (**Fig. 3**). It was found that 162 ASVs were assigned to Candidatus Brocadiales order, which were considered to be AnAOB (**Fig. S3**)^3^. Less than half of the AnAOB ASVs were assigned to the five classified AnAOB genera: *Candidatus* Kuenenia (n = 28), *Candidatus* Jettenia (n = 18), *Candidatus* Brocadia (n = 13), *Candidatus* 2-12-FULL-52-36 (n = 5), and *Candidatus* SCAELEC01 (n = 2). The results of previous study indicate that the genus *Ca.* 2-12-FULL-52-36, which has not yet been named, belongs to a novel family of AnAOB, i.e., Candidatus Bathyanammoxibiaceae^48^. *Ca.* SCAELEC01 was assigned to a novel genus of marine AnAOB distinct from the canonical *Candidatus* Scalindua genus, and that novel genus was renamed *Candidatus* Actiscalindua in the last year^49^. However, approximately 59.3% of AnAOB ASVs remained unclassified at the genus rank, indicating that there are still many unexplored AnAOB.

**Figure 3.**
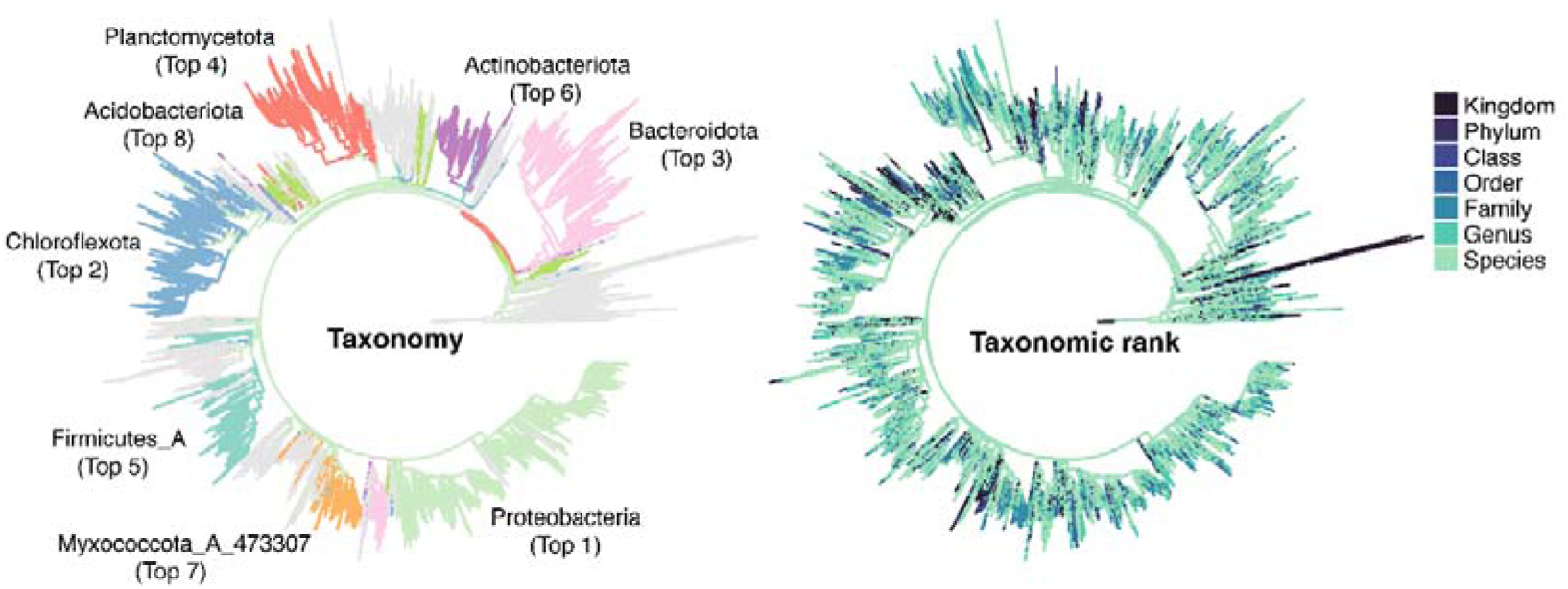
Phylogenetic tree and taxonomy of the identified ASVs. Left tree: the color of branch indicates the taxonomy of each given ASV at the phylum rank. For clarity, only the top 8 phyla were colored. Right tree: the color of branch indicates the lowest taxonomic rank of each given ASV.

Relative abundance of species was further characterized among the different types of systems. The top 5 phyla in terms of average relative abundance were, in order, Proteobacteria (28.4%), Planctomycetota (20.8%), Chloroflexota (17.8%), Bacteroidota (13.0%), and Acidobacteriota (3.0%) (**Fig. 4A**). The accumulated relative abundance the top 10 phyla in anammox communities was up to 91.4%, implying that they represent the vast majority of the community bacteria. Meanwhile, the relative abundance of these phyla spanned across systems and samples. Specifically, Proteobacteria, Planctomycetota, Chloroflexota, and Bacteroidota spanned more than 30% in abundance across biofilm, granular sludge, flocculent sludge, and planktonic cell systems. The result suggests that there is a great deal of flexibility in anammox community assembly, but that the species of its major members are restricted to the phyla described above. At the genus rank, a total of eight bacterial genera dominated the community excluding AnAOB genera and unclassified genera (**Fig. 4B**). They were assigned to three different phyla, i.e., Proteobacteria, Chloroflexota, and Bacteroidota. The accumulated relative abundance of the top 10 genera was up to 30.4%, implying that they are important components of the anammox communities.

**Figure 4.**
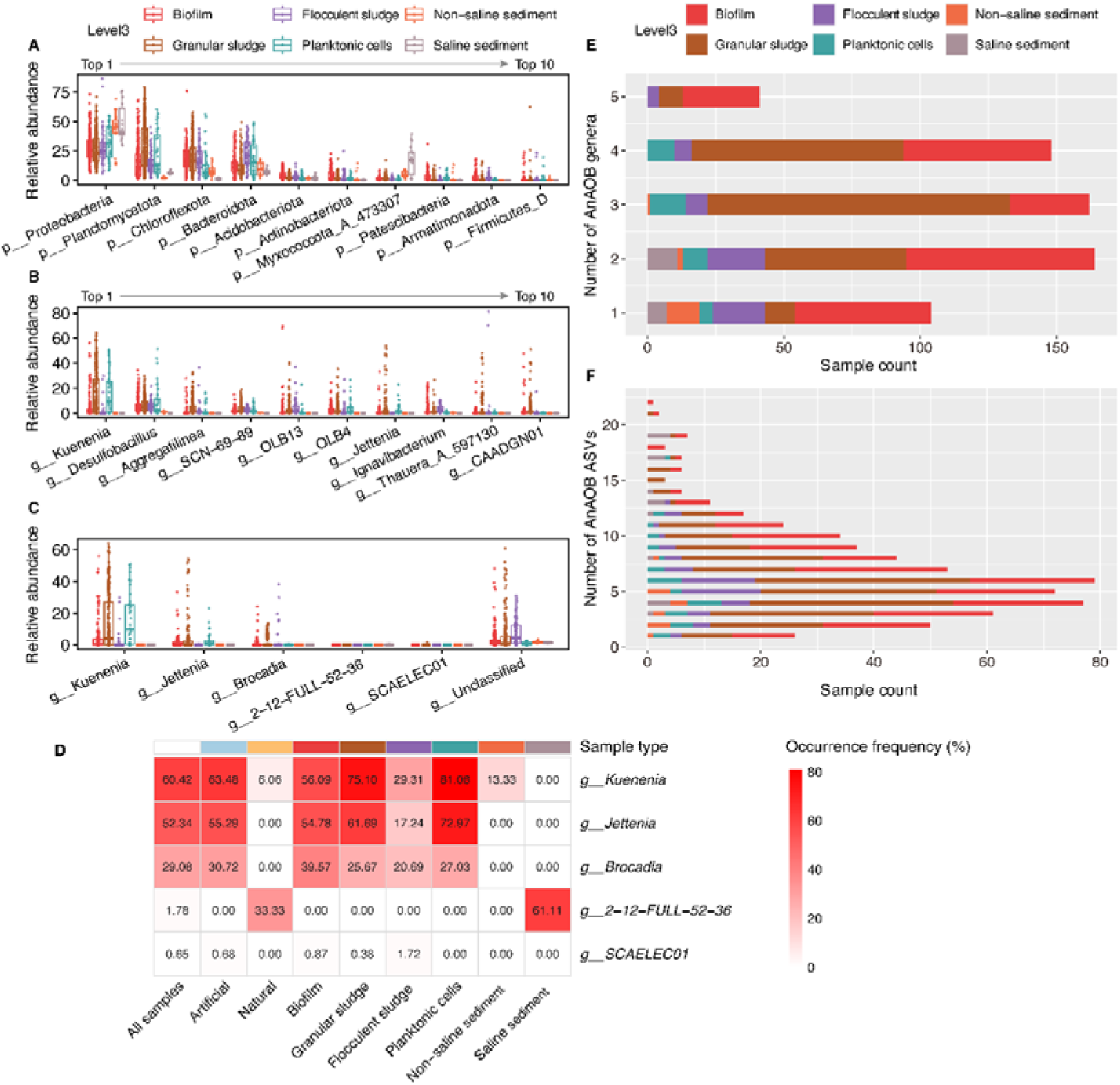
Relative abundance of taxa and number of AnAOB species in anammox communities. (**A**) Relative abundance of the top 10 phyla in different types of samples. (**B**) Relative abundance of the top 10 genera in different types of samples. (**C**) Relative abundance of identified AnAOB at the genus rank in different types of samples. (**D**) Average relative abundance of identified AnAOB at the genus rank in different types of samples. (**E**) Number of identified AnAOB genera in different types of samples. (**F**) Number of identified AnAOB ASVs in different types of samples.

*Ca.* Kuenenia, *Ca.* Jettenia, and *Ca.* Brocadia were more commonly found AnAOB among identified AnAOB genera in the samples from the artificial system, with high average relative abundances and frequency of occurrences, but spanning across systems and samples (**Fig. 4C** and **Fig. 4D**). Five AnAOB (ASV_1, ASV_20, ASV_34, ASV_367, and ASV_7) were highly adaptable in the artificial systems, with an occurrence frequency greater than 33% (**Fig. S4**). About 83% and about 95% of the samples contained two and more AnAOB genera or ASVs (**Fig. 4E** and **Fig. 4F**), and some samples even contained up to 5 AnAOB genera or 22 AnAOB ASVs. This result indicates that AnAOB tend to coexist with multiple species, i.e., one of the conditions for their survival may be the presence of genetically similar and functionally similar species.

### Core taxa and CRAT of anammox communities

To gain insight into the composition and structure of anammox communities, three types of core taxa and CRAT were identified and analyzed across all 619 samples at the genus and species rank. As shown in **Fig. 5** and **Supplementary Table 6**, 24, 34, and 31 strict, general, and loose core genera were identified, respectively. Their average relative abundance accounted for more than 60.1% and 5.8% in artificial and natural systems, respectively. Although these core genera numbered less than 2.8% of the observed genera (**Fig. 6A**), in terms of relative abundance these core genera comprised the community skeleton among artificial anammox systems (**Fig. 6B**). Besides, 360 CRAT genera related with specific culture conditions were also identified, with an average relative abundance of 10.7%. At the species rank, 16, 27 and 12 strict core, general core, and loose core species (**Fig. 5** and **Supplementary Table 7**) were identified, accounting for 1.2% of the observed species, respectively. These core species were also important components of the artificial system community, with the average relative abundance accounting for more than 35.1% (**Fig. 6C**). The number and species of core taxa are completely consistent in different artificial systems, and their relative abundances are high, which should be essential for artificial anammox systems. The results of core taxa identified in this study expand the scope of previous core taxa based on metagenomic data and provide more taxa targets for attention^8^. Meanwhile, the shared core taxa identified by the two independent studies will be the primary targets for anammox community studies (**Supplementary Table 8** and **Supplementary Table 9**), particularly the four strict core genera: *Desulfobacillus*, *Nitrosomonas*, *QUBU01*, and *OLB14*.

**Figure 5.**
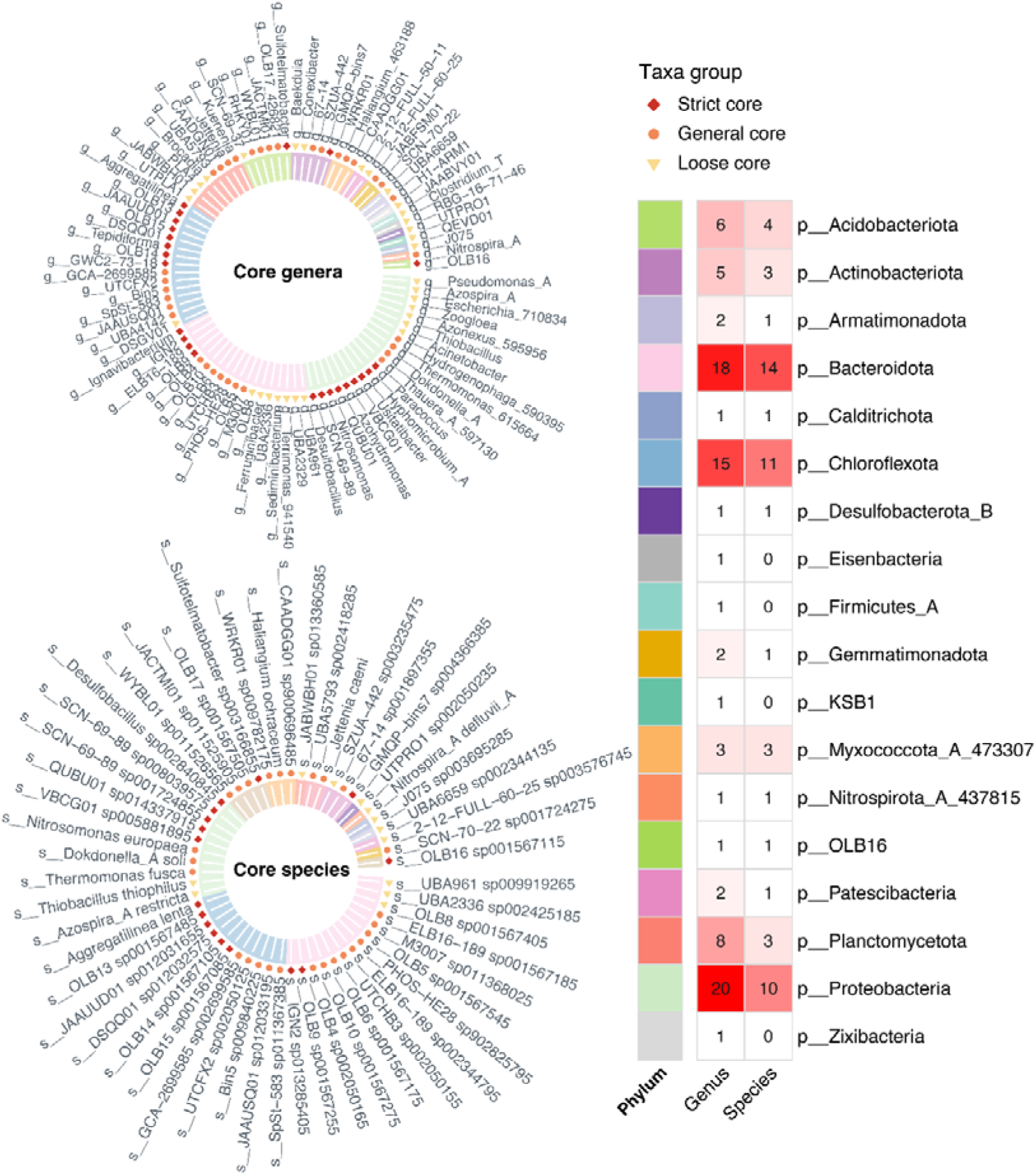
Taxonomy of core genera and species in anammox communities. In the left circular plot, the background color of each taxon corresponds to its phylum-rank classification, while distinct graphic symbols indicate the core categories (i.e., strict core, general core, and loose core). The right heatmap quantifies phylum-specific occurrences of core taxa. Numbers within heatmap cells indicate the number of core genera or species.

**Figure 6.**
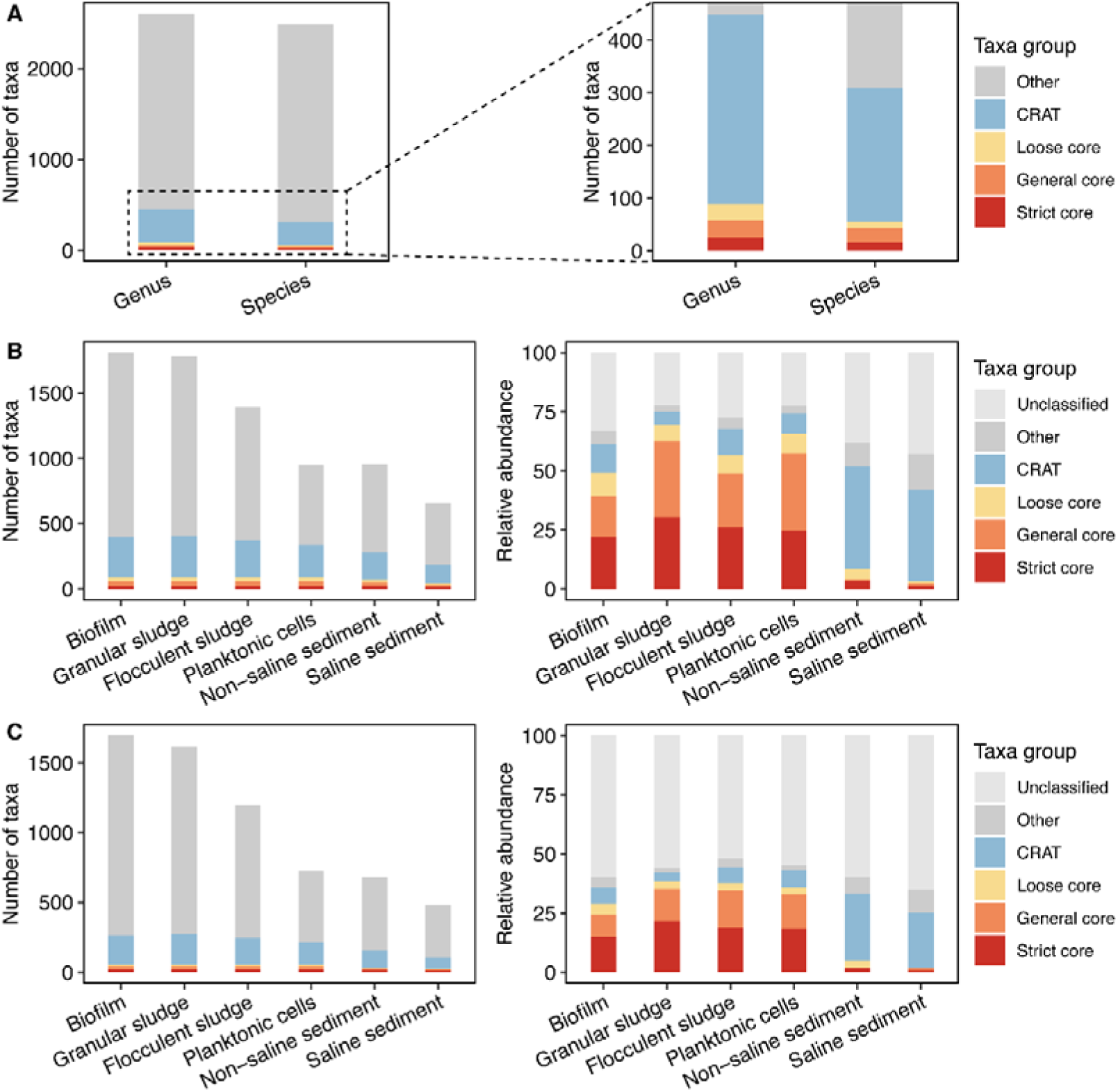
Number and abundance of core taxa and CRAT across different anammox communities. Taxa that were not assigned at the given taxonomic rank were combined as “unclassified”. Except for the five taxa (i.e., strict core, general core, loose core, CRAT, and unclassified), the rest of the taxa were combined into “other”. (**A**) Number of core taxa and CRAT identified in all samples at the genus and species rank. (**B**) Number and average relative abundance of core taxa and CRAT across the level3 samples at the genus rank. (**C**) Number and average relative abundance of core taxa and CRAT across the level3 samples at the species rank.

The core genera were mainly assigned to Proteobacteria, Chloroflexota, and Bacteroidota (**Fig. 5**). Four of the top 10 previously identified genera by average relative abundance are general core genera, and the other six were strict core genera (*Aggregatilinea*, *Desulfobacillus*, *SCN-69-89*, *OLB13*, *OLB14*, and *Ignavibacterium*). These six strict core genera all belong to microbial taxa that are difficult to culture. So far, only one species of *Aggregatilinea* has been successfully isolated, which was proved to be a slow-growing filamentous bacterium^50^. The genus *Ignavibacterium* also contains only one slow-growing pure culture (*Ignavibacterium album*), which is an organoheterotrophy bacterium with the potential to reduce nitrite to ammonium^50^. Functional descriptions of the remaining core genera were derived mainly from metagenomic analyses. *Desulfobacillus* members have sulfur compound reduction and denitrification potential^51^. Members of *OLB13* were found to contribute to nitrite accumulation, while members of *OLB14* were reported to have the complete DNRA pathway and could provide substrates for AnAOB and remove organic matter^11,52^. Members of these two genera may be involved in the process of nitrogen metabolism in anammox systems. Overall, the roles of these six strict core genera in anammox systems have not been fully explored, and difficulty in cultivation is one of the major obstacles. The evolving bacteria-specific spatial omics is a promising method to address this bottleneck^53^.

There are four strict core species all assigned to Aggregatilineales order (**Fig. S5**), and they may play a unique and indispensable role in anammox communities. The average relative abundance of strict core species spanned across different systems, which may be attributed to differences in habitat and culture conditions. Notably, *Desulfobacillus sp002840845* within Burkholderiales_597432 was the most abundant among all strictly core species, with an average relative abundance of more than 5.6% in all the artificial anammox systems. Pure cultures were not available for all of the 16 strictly core species identified, except *Aggregatilinea lenta*^50^. In the future, the function and role of these strict core species in anammox communities can be analyzed with the applying of multi-omics, especially *Aggregatilinea lenta* and *Desulfobacillus sp002840845*.

### Stable relationships between AnAOB and companion bacteria

In order to investigate the stable interactions between AnAOB and their companion bacteria, four types of artificial anammox systems (biofilm, granular sludge, flocculent sludge, and planktonic cells) were performed with independent construction of co-abundance network and analysis. All companion bacteria stably correlated with AnAOB were identified through the consistency of correlation between species across different networks. A total of 3,722 bacterial species including 28 AnAOB were found in the four types of anammox systems. Out of the possible 103,810 ASV pairs (related to AnAOB), only a very small fraction (0.2%) showed stable relationships (**Fig. S6**). Eight AnAOB were identified to be stable correlated with 147 companion bacteria (**Fig. 7** and **Supplementary Table 10**), and they form a stable co-abundance network with 208 weighted edges (**Fig. 8A** and **Supplementary Table 11**). About 16.8% of the network members had low abundance (with average relative abundance < 0.01%) in artificial anammox systems, which could be easily ignored by conventional metagenomics. In this network, the number of positive correlations (76.4%) was much larger than the number of negative correlations (23.6%), with 105 and 35 species positively and negatively correlated with AnAOB, respectively. Seven additional species were two-sided, with positive correlations for some AnAOB and negative correlations for others. Significant correlations (P < 0.01) were found between these species under both attached (biofilm or granular sludge) and suspended (flocculent sludge or planktonic cells) growth modes, indicating that their interactions span different growth modes.

**Figure 7.**
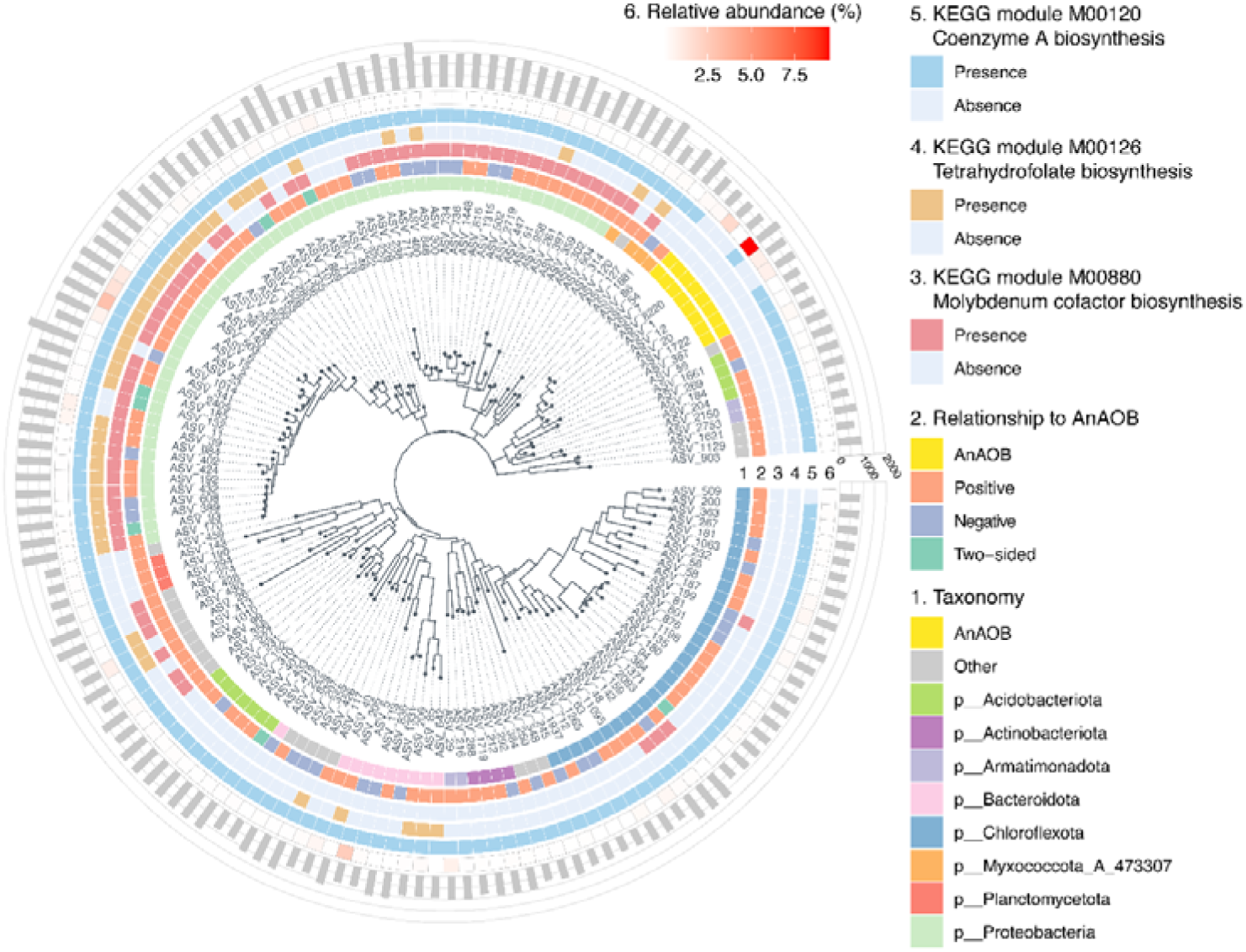
Phylogenetic tree of AnAOB and stably correlated companion bacteria in artificial anammox systems. There are six circles of annotation information outside the phylogenetic tree, which are taxonomy at the phylum rank, relationship to AnAOB, molybdenum cofactor biosynthesis potential, tetrahydrofolate biosynthesis potential, coenzyme A biosynthesis potential, and average relative abundance (within artificial anammox systems) of each given bacterium from the inner circle to the outer circle. Phyla containing fewer than four ASVs were combined into “other”. The bar represents the number of predicted KEGG orthology (KO) for each given ASV.

**Figure 8.**
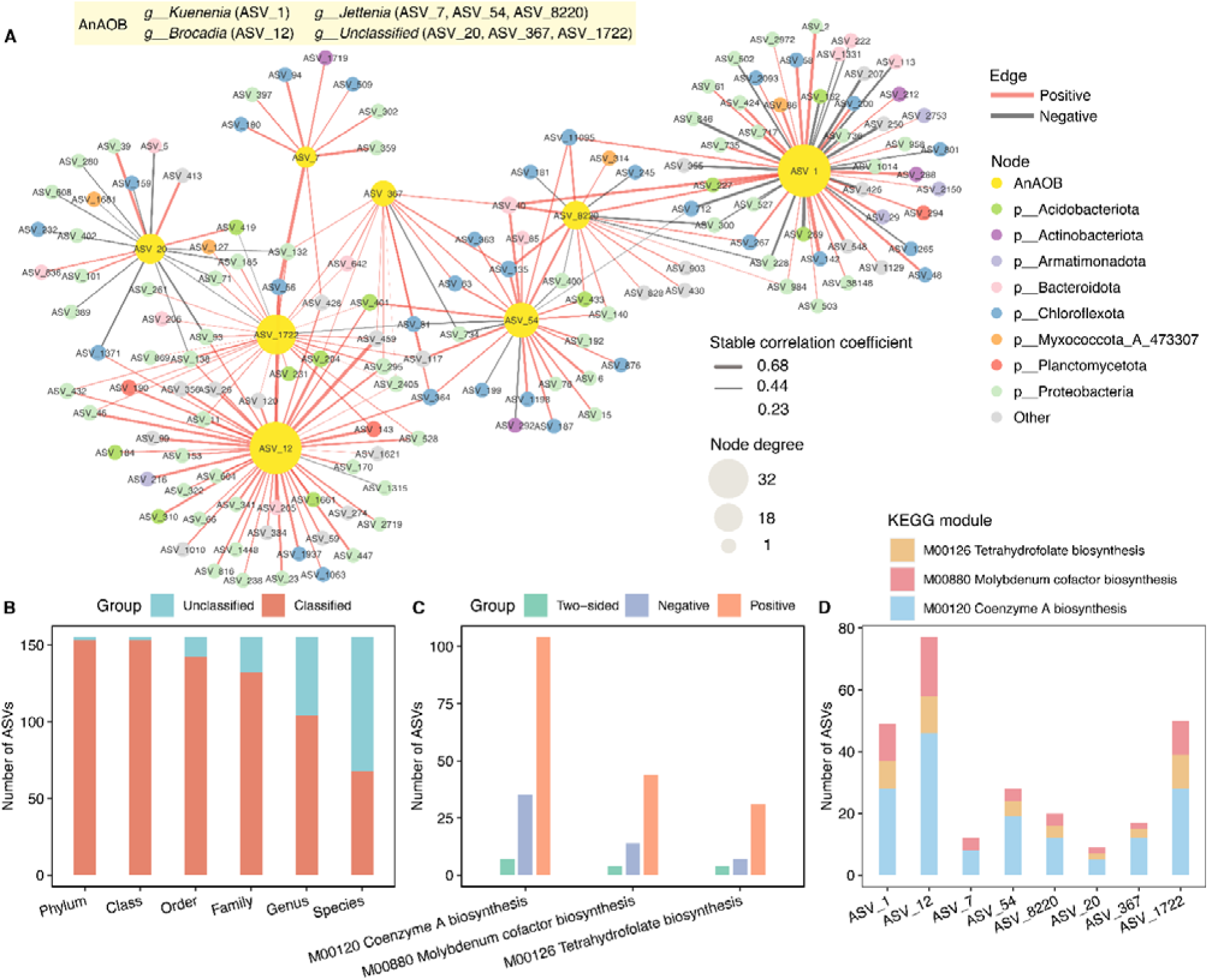
Stable relationships of AnAOB in artificial anammox systems (biofilms, granular sludges, flocculent sludges, and planktonic cells). (**A**) Stable co-abundance network of AnAOB and companion bacteria. The size of the node indicates the degree, and the color of the node indicates taxonomy at the phylum rank. The color and thickness of the edges indicate the positive and negative and the magnitude of the stable correlation coefficient. (**B**) Number of classified and unclassified ASVs at different taxonomic ranks that make up the stable co-abundance network. (**C**) Number and group of companion bacteria with potential for molybdenum cofactor, tetrahydrofolate, and coenzyme A synthesis. Positive, negative and two-sided species were not double-counted. (**D**) Number and group of positively correlated companion bacteria with potential to biosynthesis molybdenum cofactor, tetrahydrofolate biosynthesis, and coenzyme A. Some companion bacteria had two or three co-factor synthesis potentials at the same time, and therefore were counted in duplicate at the time of statistics.

The stable co-abundance network contained members of 21 known phyla and 78 known genera (**Fig. 8B**), with more than 67.1% of these species classified at the genus rank and above. The stable companion bacteria of AnAOB were mainly Proteobacteria (n = 59) and Chloroflexota (n = 30) members. It is noteworthy that the vast majority (96%) of known genera contained only one to two stable companion bacteria, whereas *DSQQ01* and *Desulfobacillus* contained six and four stable companion bacteria, respectively. This result again suggests that members of strict core genera *DSQQ01* and *Desulfobacillus* play key roles in the anammox systems. Five of the eight AnAOB with stable relationship were assigned to classified genera, *Ca.* Kuenenia (n = 1), *Ca.* Brocadia (n = 1), *Ca.* Jettenia (n = 3), and the other three were unclassified at the genus rank. The species and number of stable companion bacteria varied greatly among the different AnAOB genera or species, but at the same time there were a small number of shared stable companion bacteria (**Fig. S7**). Some companion bacteria (e.g., ASV_428, ASV_56, and ASV_364) could interact with more than one AnAOB, and some AnAOB pairs (e.g., ASV_1 and ASV_8220, ASV_12 and ASV_1722) were stably and positively correlated with each other (**Fig. 8A**).

Previous studies have shown that AnAOB performs CO_2_ fixation via the Wood-Ljungdahl pathway^54^. Molybdenum cofactor and tetrahydrofolate play an important role in this pathway^55,56^. However, the functional prediction results showed that all the AnAOB did not have the complete molybdenum cofactor and tetrahydrofolate biosynthesis pathways (**Fig. S8**), suggesting that these AnAOB were required to obtain essential cofactors from other members within communities. Among all stable companion bacteria, 62 and 42 species had molybdenum cofactor and tetrahydrofolate biosynthesis potential, respectively (**Fig. 8C**). Notably, 44 and 31 species were positively correlated to AnAOB, respectively. Moreover, each of AnAOB except ASV_7 had at least one positively correlated bacterium that could biosynthesize molybdenum cofactor or tetrahydrofolate (**Fig. 8D**). This result indicates that the positively correlated bacteria identified are likely to provide molybdenum cofactor and tetrahydrofolate to AnAOB, which lends strong evidence to the conclusions obtained from previous independent and single-system studies^10,11,22^. Meanwhile, our result suggests that the above conclusion may be general in artificial anammox systems. Only three AnAOB could biosynthesize de novo coenzyme A, an important substance involved in tricarboxylic acid cycle and fatty acid biosynthesis^57^. It is well known that fatty acids, as key components of cell membranes, are essential for bacterial survival^58^. This indicates that the other five AnAOB may require other companion bacteria to provide coenzyme A. This conclusion is supported by the fact that each of AnAOB had lots of positively correlated bacteria that could biosynthesize coenzyme A (**Fig. 8D**). More than 92% of the positively correlated bacteria had no known complete microbial CO_2_ fixation pathway, indicating that most of them were heterotrophic and required exogenous organic carbon sources. As speculated by previous studies^59,60^,organic carbon sources, e.g., extracellular polymeric substance (EPS), peptides or detritus, produced by autotrophic AnAOB may support the growth of these heterotrophic bacteria.

Further attention is paid to the strong correlation in the network, that is, the stable correlation coefficient is ≥ 0.5 or ≤ −0.5. These strongly correlated species are assumed to interact more closely than others. Among the strong positively correlated bacteria of *Ca.* Kuenenia (ASV_1), it is notable that *Desulfobacillus sp002840845* (ASV_2), a strict core species in anammox communities, is highly abundant and ubiquitous. *Desulfobacillus sp002840845* had the potential to biosynthesize molybdenum cofactor and tetrahydrofolate with complete DNRA and denitrification pathways. The mutual symbiosis of *Ca.* Kuenenia and *Desulfobacillus sp002840845* should be further investigated. Eleven strong positively correlated bacteria were identified for each of *Ca.* Brocadia (ASV_12) and *Ca.* Jettenia (ASV_7, ASV_54, and ASV_8220). Notably, ASV_56 (*JAAUUD01*) showed strong positive correlations with both ASV_12 and ASV_7. Members of *Ca.* Brocadia and *Ca.* Jettenia were found to be unable to biosynthesize molybdenum cofactors and tetrahydrofolate, as well as coenzyme A. ASV_56 was able to provide molybdenum cofactor, tetrahydrofolate and coenzyme A to *Ca.* Brocadia and *Ca.* Jettenia, which adds new evidence to the previous speculation^11^.

Bioaugmentation with exogenous cofactors essential for AnAOB has been demonstrated to enhance nitrogen removal efficiency in anammox systems. For instance, folate supplementation at 8 mg/L resulted in a 67.1% decrease in the anammox granular floatation potential, which maintained the high nitrogen removal efficiency of anammox system^61^. However, in engineered anammox systems, the external addition of expensive cofactors will increase treatment costs. It may be an alternative strategy to enhance anammox system performance by regulating these heterotrophic bacteria in mutual symbiosis with AnAOB, as they can both provide cofactors and remove the constantly produced organic matter within such systems. The numerous positive heterotrophic companion bacteria identified in this study provide the regulating targets to choose from. Furthermore, in recent years, synthetic microbial communities have received much attention due to their advantages of strong environmental adaptability, high efficiency, and stability^62,63^. Analytical findings on core species identification and their stable interspecific relationships offer valuable references for synthesizing anammox communities via the top-down strategy.

Strong negative correlations of species pairs are often indicative of competition^64^. Eleven strong negative correlations were identified in the current study, involving *Ca.* Kuenenia and 11 other bacteria (**Supplementary Table 11**). Most of these companion bacteria (except ASV_846 and ASV_1014) lacked the ability to biosynthesize molybdenum cofactor or tetrahydrofolate and key genes for ammoxidation, denitrification, and DNRA, suggesting that they may compete with *Ca.* Kuenenia for essential cofactors. Both ASV_846 and ASV_1014 had DNRA and ammoxidation potential, respectively, indicating that they may compete with *Ca.* Kuenenia for substrate. These 11 bacteria could be the negative biomarkers of engineering anammox systems dominated by *Ca.* Kuenenia, that is, their appearance or overgrowth predicts a decrease in anammox activity.

### Limited stable relationships between AnAOB and companion bacteria

Bacteria are generally tightly bound together in the anammox attachment-growth systems (biofilm and granular sludge) due to the abundant and sticky EPS^65^. The anammox attachment-growth systems have numerous advantages, e.g., stronger biomass retention capacity, higher nitrogen loading rate, and higher resistance to environmental stress, compared with the anammox suspended growth system, and such systems are suitable for engineering applications^17,66^. Therefore, the stable relationships in the anammox attachment-growth systems, designated as limited stable relationships, were further explored. A total of 291 bacterial species, including three AnAOB, were found to be shared in the above four types of anammox systems.

Approximately 10.6% of the 867 possible ASV pairs exhibited limited stable relationships (**Fig. S9**). Three AnAOB (ASV_1, ASV_12, and ASV_367) were identified to be limited stable correlated with 79 companion bacteria in the anammox attachment-growth systems (**Fig. 9A** and **Supplementary Table 12**), and they formed a co-abundance network with 92 weighted edges (**Supplementary Table 13**). All 82 bacteria were assigned to 14 phyla with Proteobacteria (n = 35) and Chloroflexota (n = 14) having the most members, and more than 58.5% of the bacteria were classified at the genus rank (**Fig. 9B**). Forty-three known genera were identified, including previously identified strict core genera: *Desulfobacillus*, *OLB15*, *SCN-69-89*, *Nitrosomonas*, etc. One ASV each was assigned to *Ca.* Kuenenia, *Ca.* Brocadia, and genus unclassified AnAOB. The species and number of limited-stable-correlated bacteria varied among AnAOB (**Fig. 9A** and **Fig. 9C**).

**Figure 9.**
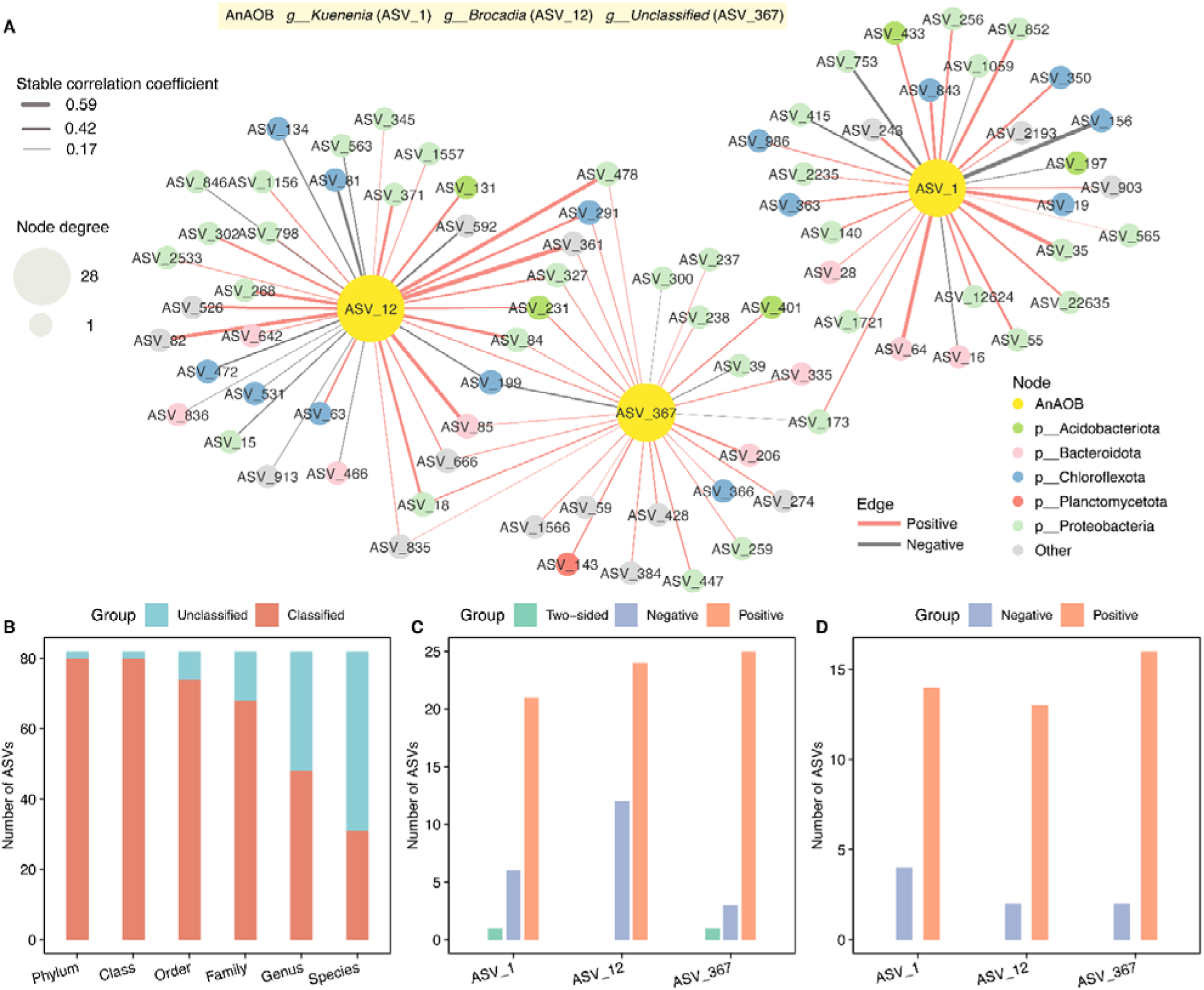
Limited stable relationships of AnAOB in anammox attachment-growth systems (biofilms and granular sludges). (**A**) Limited stable co-abundance network of AnAOB and companion bacteria. The size of the node indicates the degree, and the color of the node indicates taxonomy at the phylum rank. The color and thickness of the edges indicate the positive and negative and the magnitude of the stable correlation coefficient. (**B**) Number of classified and unclassified ASVs at different taxonomic ranks that make up the limited stable co-abundance network. (**C**) Number and group of limited stable companion bacteria per AnAOB. (**D**) Number and group of limited stable companion bacteria with interspecies electron transport potential per AnAOB.

The positive interaction between two bacteria occurs only at a tightly-bound mode, i.e., attached growth mode. One possibility is that they perform direct interspecies electron transfer (DIET)^67^. DIET is typically mediated by electroactive bacteria that perform extracellular electron transport (EET) through their own conductive structures, e.g., conductive type pili (T4P) and cytochrome c^68^. Previous studies have shown that AnAOB (e.g., *Ca.* Kuenenia, *Ca.* Brocadia, and *Ca.* Scalindua) are electroactive and can carry out EET, transferring the electrons obtained from oxidizing ammonium to the extracellular electron acceptor to generate energy^69,70^. T4P have been found to play key roles in bacterial biofilm formation^71^. Therefore, it is speculated that AnAOB recruits companion bacteria with T4P to form symbiotic aggregates. The addition of such beneficial companion bacteria to engineered anammox systems may accelerate the formation of anammox biofilm or granular sludge and enhance anammox activity. In the current study, not only three AnAOB (ASV_1, ASV_12, and ASV_367), but also 42 bacteria of limited network had the potential for DIET (harboring genes encoding T4P), of which the vast majority (> 85.7%) were positively correlated bacteria of AnAOB (**Fig. S10**). Since KO numbers were not assigned to EET cytochrome c in the KEGG database, cytochrome c-mediated DIET is not discussed here. Each of AnAOB had at least 13 positively correlated bacteria with DIET potential (**Fig. 9D**). It is likely that these companion bacteria engage in mutualism with AnAOB through DIET. Furthermore, six strong positive relationships were identified in the limited network (**Supplementary Table 13**), involving two AnAOB (*Ca.* Kuenenia and *Ca.* Brocadia) and other six bacteria (ASV_35, ASV_64, ASV_82, ASV_85, ASV_361, and ASV_478). At present, the role of T4P in anammox attachment-growth systems are underexplored and should be further delved into. The positively correlated bacteria, particularly these strongly correlated bacteria, could be the primary targets for future studies on tightly-bound interactions between AnAOB and their companion bacteria. Moreover, analytical findings on limited stable relationships can also provide guidance for the synthesis of anammox biofilm or granular sludge communities, such engineered synthetic anammox communities will be robust and efficient.

## Conclusions

In summary, we systematically analyzed bacterial diversity, core taxa, and (limited) stable correlations of AnAOB based on 619 global anammox samples. In artificial anammox systems, *Ca.* Kuenenia, *Ca.* Jettenia, and *Ca.* Brocadia are common AnAOB, and different AnAOB species coexist in one system. Different artificial anammox systems share a number of core taxa that they can form the backbone of bacterial communities. Co-abundance network analysis across different artificial anammox systems showed that AnAOB have some stable or limited stable relationships with companion bacteria. Stable positively-correlated bacteria may provide essential cofactors (e.g., molybdenum cofactor, tetrahydrofolate, and coenzyme A) to AnAOB. Limited stable positively-correlated bacteria may interact with AnAOB through T4P within anammox attachment-growth systems. This work advances the understanding of anammox communities, anammox core microbiome, and AnAOB symbiotic relationships from a global-scale perspective. The (limited) stable companion bacteria identified in this study can be used as regulatory targets for bioaugmentation or biomarkers of deterioration for engineered anammox systems. Although deciphering the symbiotic mechanisms of anammox communities in different forms of sludge faces many difficulties, the ongoing microscopic and mesoscopic spatial omics is likely to overcome these limitations.

## Data availability

The reads of 94 anammox sludge samples collected in this study have been deposited in the China National Center for Bioinformation (accession number: PRJCA034550). The reads of other 525 samples are available in the NCBI SRA database via the accession number provided in **Supplementary Table 2.** The 16S rRNA amplicon dataset of anammox communities is available on Figshare (DOI: 10.6084/m9.figshare.28570991).

## Acknowledgements

This work was financially supported by the National Natural Science Foundation of China (52370030 and 52400152) and Chongqing Entrepreneurship and Innovation Support Program for Returned Overseas Scholars (cx2024032).

## Author contributions

Y.-P.C. and Y.-C.W. conceived and designed the study. Y.-C.W. performed bioinformatics analysis and visualization. Y.-C.W., H.-M.F., H.-Y.L., H.W., Z.-H.Z., J.-H.H., and L.Z. contributed to sample and metadata collection. Y.-C.W. and H.-M.F. wrote the manuscript. Y.-P.C.., L.Z., P.Y., and M.-J.G. revised the manuscript.

