## Supplementary information for "Microbial community characteristics and stable relationships centered on anammox bacteria revealed by global-scale analysis"

This supplementary information contains 13 pages, including 10 figures and this cover page.

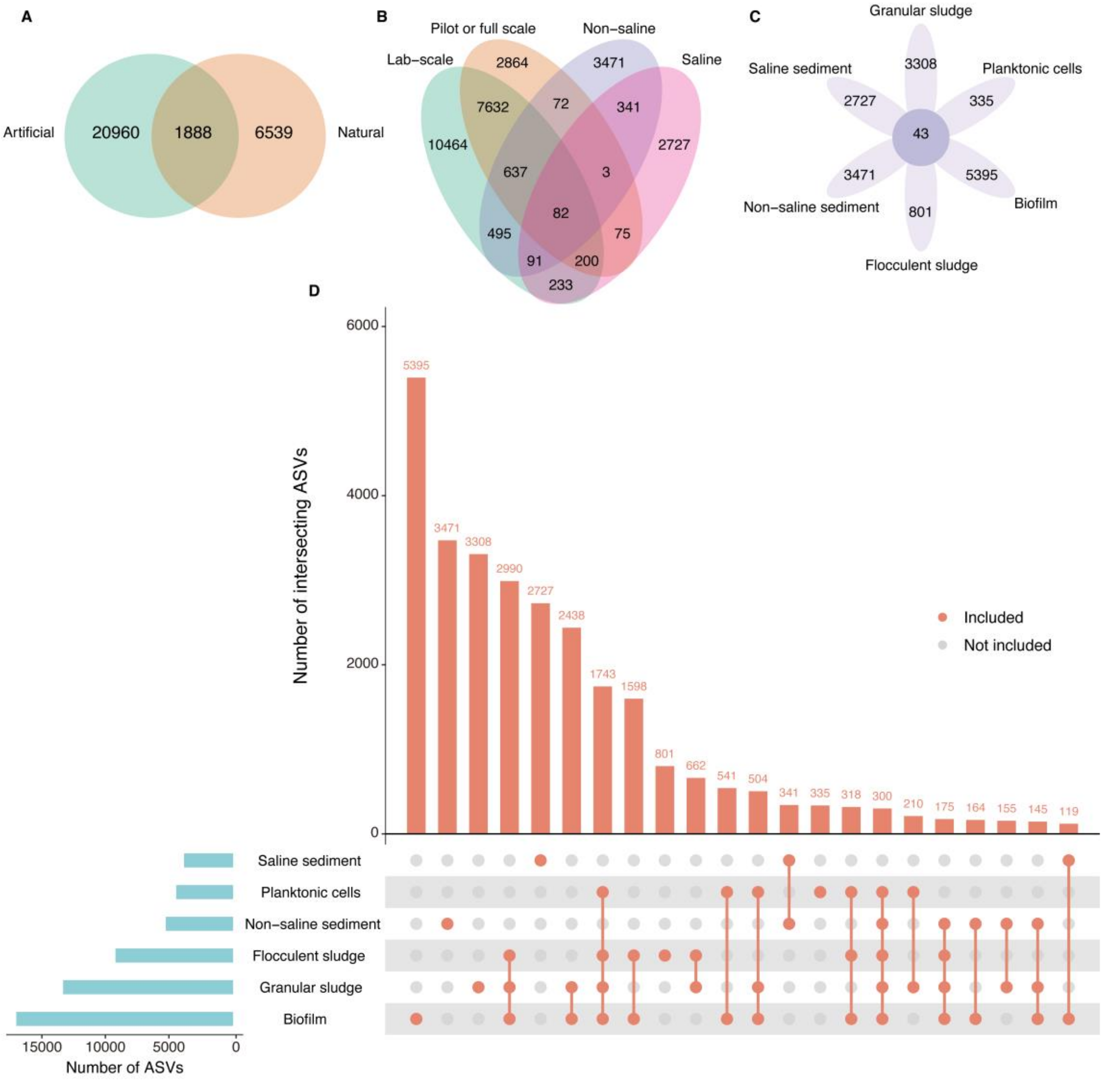


**Figure S1.** Species intersection size for different anammox communities. (**A**) Species intersection for level1 samples. (**B**) Species intersection for level2 samples. (**C**) Species intersection for level3 samples. The number in the petal indicates the number of species unique to samples of each given type, and the number in the center of the petal indicates the number of species common to all level3 samples. (**D**) Upset plot of the species intersection size for level3 samples. The bar in the upper right corner represents the number of species that are intersection or unique, and the bar in the lower left corner represents the number of all species identified for each given type of samples. For clarity, only intersections with a number greater than 100 are shown.

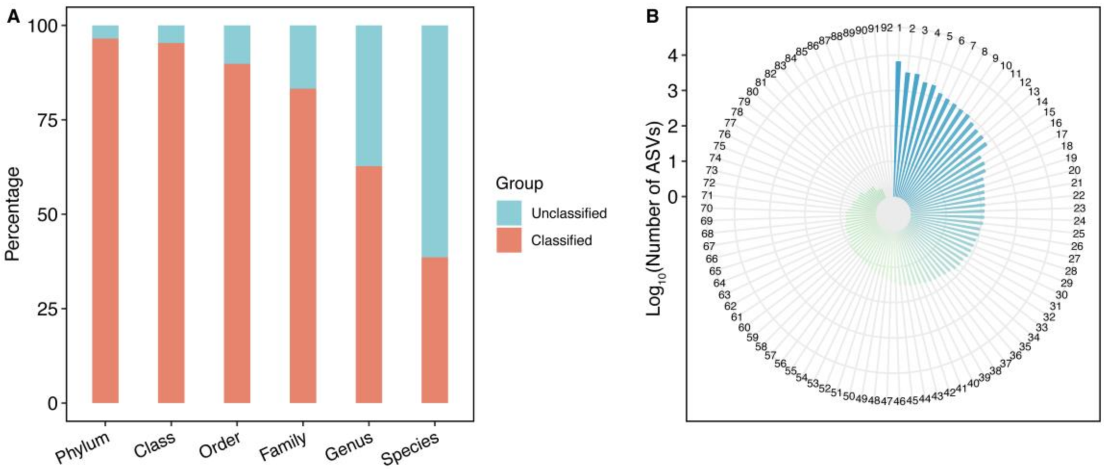


**Figure S2.** Proportion of classification and number of ASVs identified. (**A**) Percentage of classification for identified ASVs at the phylum, class, order, family, genus and species rank. (**B**) The number of ASVs identified at the phylum rank, the numbers 1 to 92 in the figure represent the phylum name codes, the specific information is shown in the **Supplementary Table 5**.

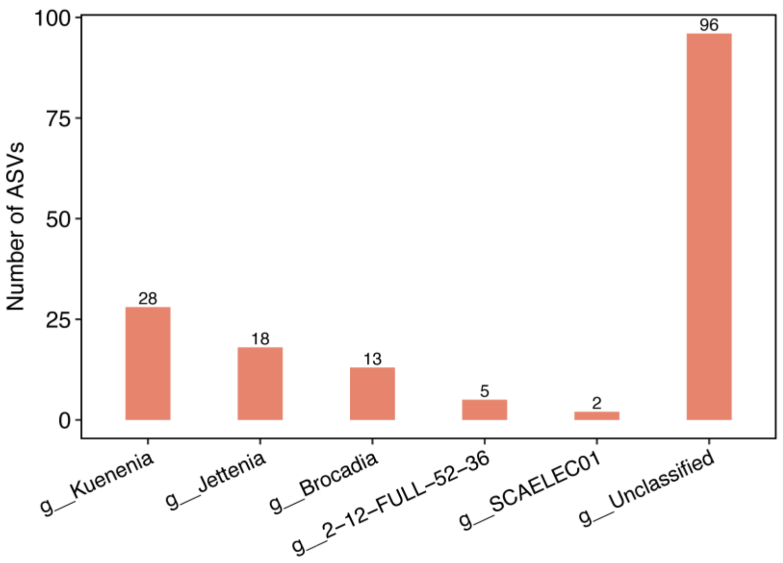


**Figure S3.** Taxonomy and number of identified AnAOB ASVs.

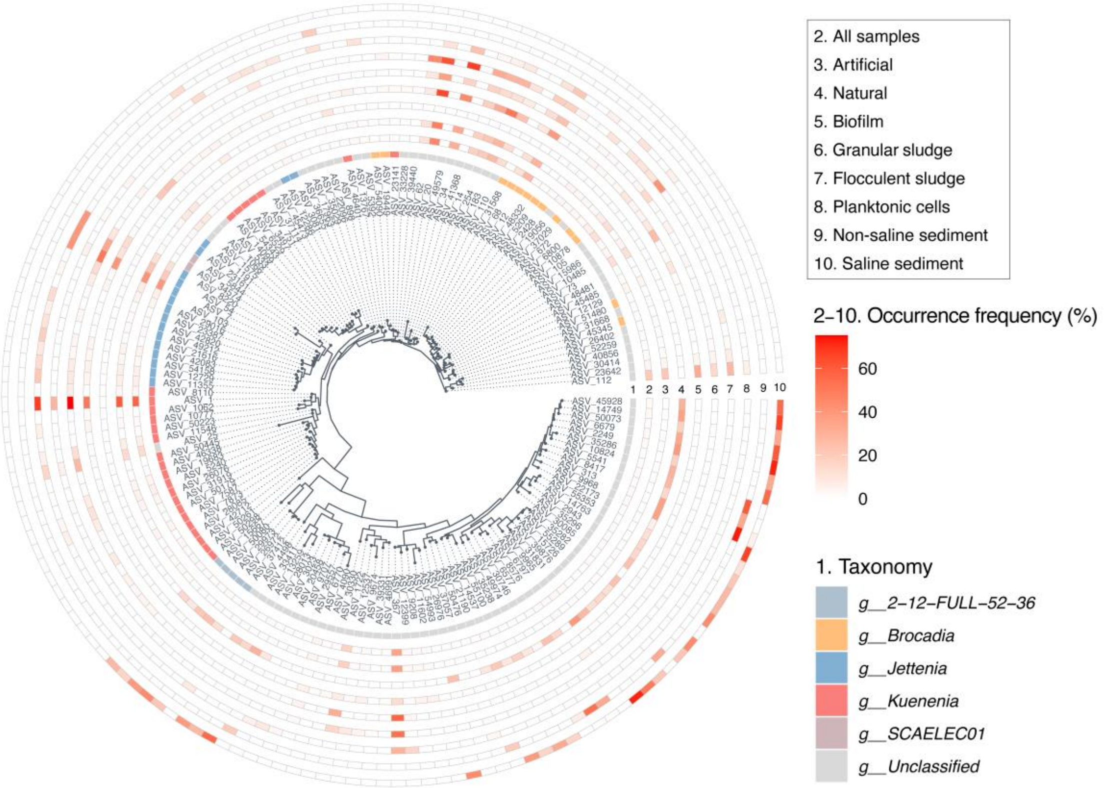


**Figure S4.** Phylogenetic tree of identified AnAOB and their occurrence frequency across different types of samples. The heat map colors at tracks 2 to 10 on the outer side of the tree represent the occurrence frequency of each given AnAOB. To make the occurrence frequency of AnAOB comparable, the statistics after rarefaction present here. Three AnAOB ASVs were absence in rarefied ASV table.

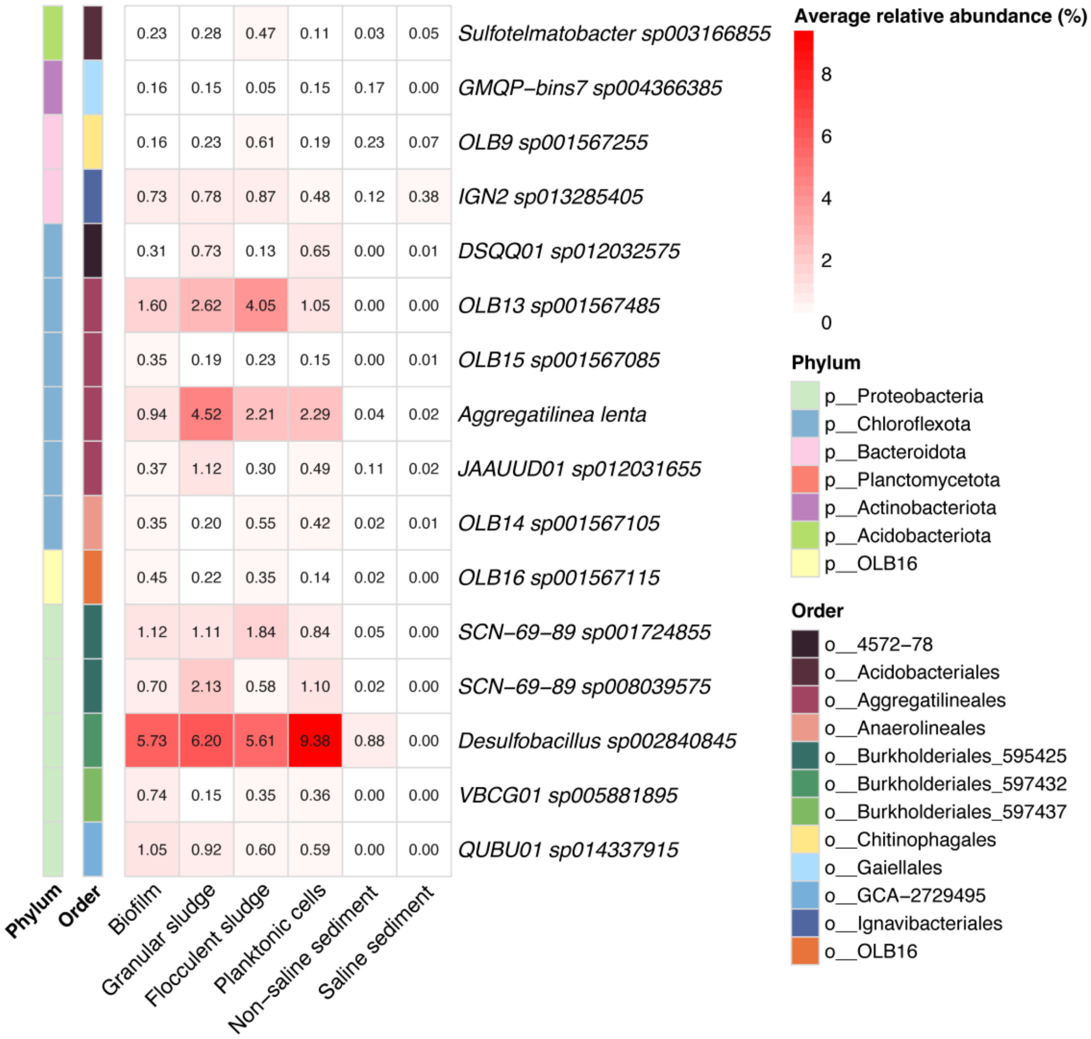


**Figure S5.** Average relative abundance of 16 strict core species across the level3 samples.

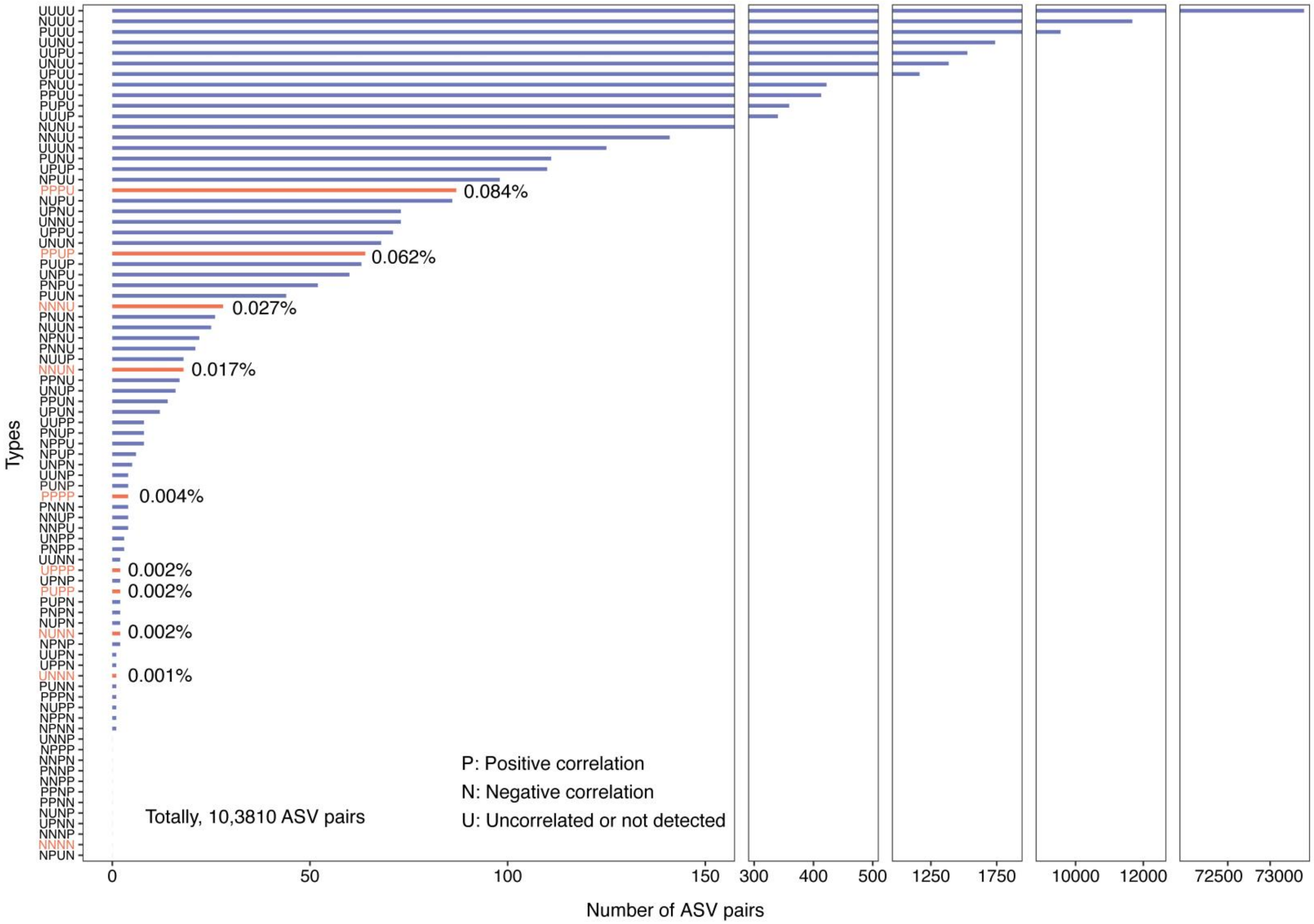

Types
UUUU
NUUU
PUUU
UUNU
UUPU
UNUU
UPUU
PNUU
PPUU
PUPU
UUUP
NUNU
NNUU
UUUN
PUNU
UPUP
NPUU
PPPU
NUPU
UPNU
UNNU
UPPU
UNUN
PPUP
PUUP
UNPU
PNPU
PUUN
NNNU
PNUN
NUUN
NPNU
PNNU
NUUP
NNUN
PPNU
UNUP
PPUN
UPUN
UUPP
PNUP
NPPU
NPUP
UNPN
UUNP
PUNP
PPPP
PNNN
NNUP
NNPU
UNPP
PNPP
UUNN
UPPP
UPNP
PUPP
PUPN
PNPN
NUPN
NUNN
NPNP
UUPN
UPPN
UNNN
PUNN
PPPN
NUPP
NPPN
NPNN
UNNP
NPPP
NNPN
PNNP
NNPP
PPNP
PPNN
NUNP
UPNN
NNNP
NNNN
NPUN
0.084%
0.062%
0.027%
0.017%
0.004%
0.002%
0.002%
0.002%
0.001%
P: Positive correlation
N: Negative correlation
U: Uncorrelated or not detected
Totally, 10,3810 ASV pairs
0
50
100
150
300
400
500
1250
1750
10000
12000
72500
73000
Number of ASV pairs

**Figure S6.** The number of ASV pairs (related to AnAOB) identified in artificial anammox systems (biofilms, granular sludges, flocculent sludges, and planktonic cells). The three letters show the correlations (P for positive, N for negative, and U for uncorrelated or not detected) in order of the ASV pairs in biofilms, granular sludges, flocculent sludges, and planktonic cells. Stable correlations across anammox systems (NNNN, PPPP, UNNN, NUNN, NNUN, NNNU, UPPP, PUPP, PPUP, and PPPU) were shown in light red, with percentages also marked.

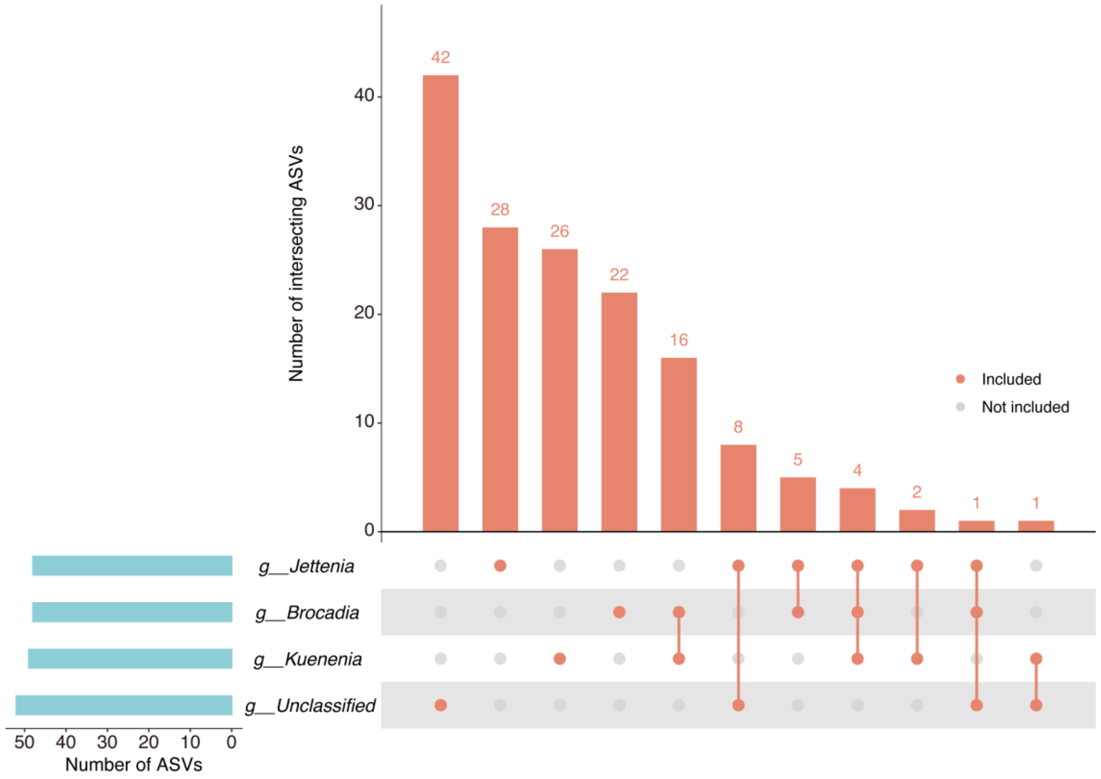


**Figure S7.** Upset plot of stable correlated companion bacteria for different AnAOB genera. The bar in the upper right corner represents the number of companion bacteria that are intersection or unique, and the bar in the lower left corner represents the number of all companion bacteria identified for each given AnAOB genus. All intersections are shown.

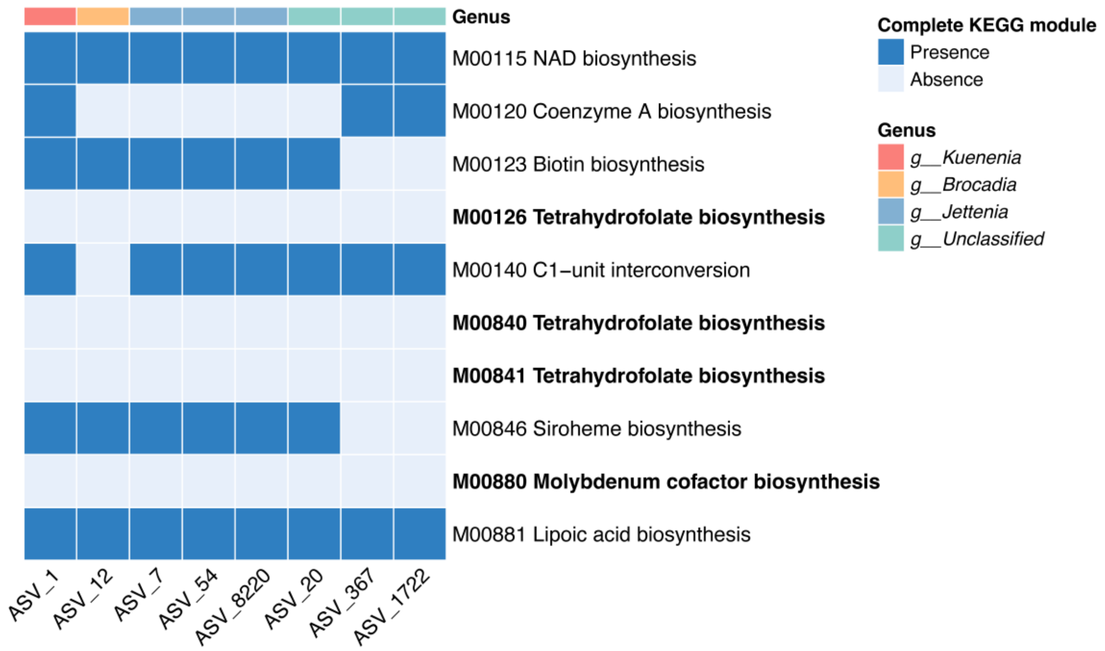


**Figure S8.** Predicted cofactor and vitamin metabolism pathways of AnAOB.

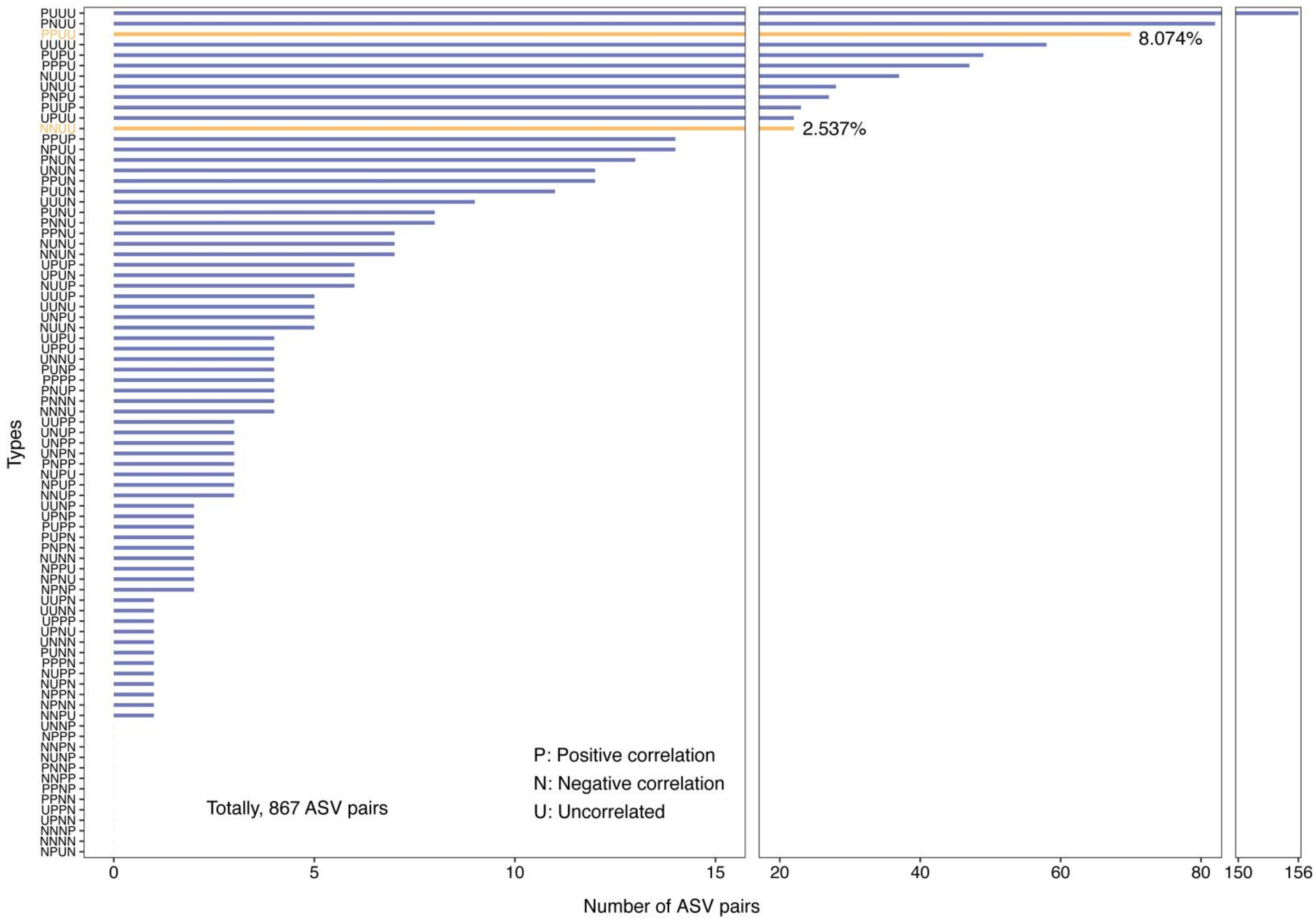

Types
PUUU
PNUU
PPUU
UUUU
PUPU
PPPU
NUUU
UNUU
PNPU
PUUP
UPUU
NNUU
PPUP
NPUU
PNUN
UNUN
PPUN
PUUN
UUUN
PUNU
PNNU
PPNU
NUNU
NNUN
UPUP
UPUN
NUUP
UUUP
UUNU
UNPU
NUUN
UUPU
UPPU
UNNU
PUNP
PPPP
PNUP
PNNN
NNNU
UUPP
UNUP
UNPP
UNPN
PNPP
NUPU
NPUP
NNUP
UUNP
UPNP
PUPP
PUPN
PNPN
NUNN
NPPU
NPNU
NPNP
UUPN
UUNN
UPPP
UPNU
UNNN
PUNN
PPPN
NUPP
NUPN
NPPN
NPNN
NNPU
UNNP
NPPP
NNPN
NUNP
PNNP
NNPP
PPNP
PPNN
UPPN
UPNN
NNNP
NNNN
NPUN
8.074%
2.537%
P: Positive correlation
N: Negative correlation
U: Uncorrelated
Totally, 867 ASV pairs
0
5
10
15
20
40
60
80
150
156
Number of ASV pairs

**Figure S9.** The number of shared ASV pairs (related to AnAOB) identified in artificial anammox systems (biofilms, granular sludges, flocculent sludges, and planktonic cells). The three letters show the correlations (P for positive, N for negative, and U for uncorrelated or not detected) in order of the ASV pairs in biofilms, granular sludges, flocculent sludges, and planktonic cells. Limited stable correlations across anammox systems (PPUU and NNUU) were shown in yellow, with percentages also marked.

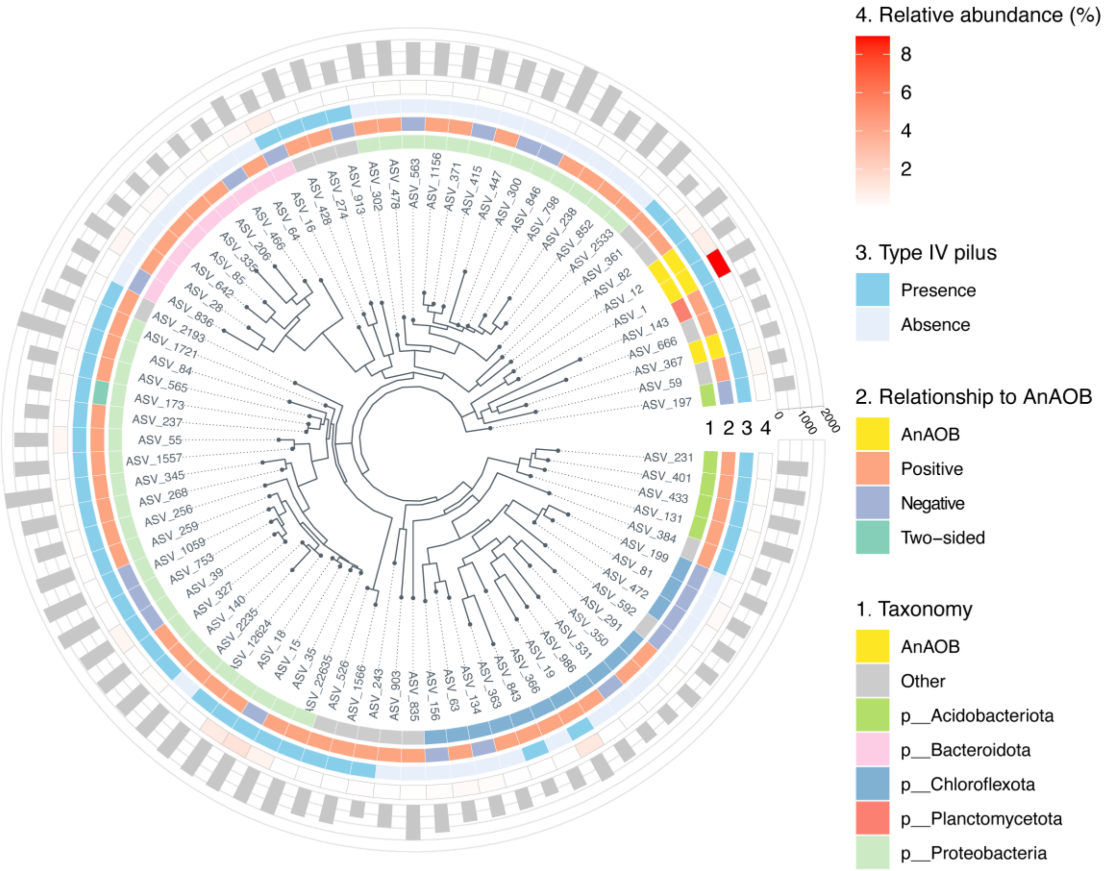


**Figure S10.** Phylogenetic tree of AnAOB and limited stably-correlated companion bacteria. There are four circles of annotation information outside the phylogenetic tree, which are taxonomy at the phylum rank, relationship to AnAOB, type IV pilus assembly potential, and average relative abundance (within artificial anammox systems) of each given bacterium from the inner circle to the outer circle. Phyla containing fewer than four ASVs were combined into “other”. The bar represents the number of predicted KEGG orthology (KO) for each given ASV.
